# GABA-mediated tonic inhibition differentially modulates gain in functional subtypes of cortical interneurons

**DOI:** 10.1101/613075

**Authors:** Alexander Bryson, Robert Hatch, Bas-Jan Zandt, Christian Rossert, Samuel Berkovic, Christopher A Reid, David B Grayden, Sean Hill, Steven Petrou

## Abstract

The binding of GABA to extra-synaptic GABA_A_ receptors generates tonic inhibition that acts as a powerful modulator of cortical network activity. Despite GABA being present at low levels throughout the extracellular space of the brain, previous work has shown that GABA may differentially modulate the excitability of neuron subtypes through variation in chloride gradient. Here, we introduce a distinct mechanism through which extracellular GABA can differentially modulate the excitability of neuron subtypes through variation in neuronal electrophysiological properties. Using biophysically-detailed computational models, we found that tonic inhibition enhanced the responsiveness (or gain) of models with electrophysiological features typically observed in somatostatin (Sst) interneurons and reduced gain in models with features typical for parvalbumin (Pv) interneurons. These predictions were experimentally verified using patch-clamp recordings. Further analysis revealed that differential gain modulation is also dependent upon the extent of outward rectification of the GABA_A_ receptor-mediated tonic current. Our detailed neuron models demonstrate two subcellular consequences of tonic inhibition. First, tonic inhibition enhances somatic action potential repolarisation by increasing current flow into the dendritic compartment. This enhanced repolarisation then reduces voltage-dependent potassium currents at the soma during the afterhyperpolarisation. Finally, we show that reductions of potassium current selectively increase gain within neurons exhibiting action potential dynamics typical for Sst interneurons. Potassium currents in Pv-type interneurons are not sensitive to this mechanism as they deactivate rapidly and are unavailable for further modulation. These findings introduce a neuromodulatory paradigm in which GABA can induce a state of differential interneuron excitability through differences in intrinsic electrophysiological properties.

## INTRODUCTION

γ-aminobutyric acid (GABA) is the predominant inhibitory neurotransmitter of the mammalian brain and regulates neuronal excitability through two modes of transmission: phasic and tonic inhibition(1). Phasic inhibition is mediated by presynaptic GABA release that activates GABA_A_ and GABA_B_ receptors within the post and peri-synaptic membrane. Tonic inhibition is mediated by extracellular, or ‘ambient’, GABA that diffuses throughout the extracellular space and activates GABA_A_ receptors within the extra-synaptic membrane(2). Tonic inhibition exerts a powerful neuromodulatory influence across most brain regions including the cerebral cortex, hippocampus, cerebellum and thalamus(3).

Extra-synaptic GABA_A_ receptors that mediate tonic inhibition possess unique subunit composition, electrophysiologic and pharmacologic properties compared to postsynaptic GABA_A_ receptors(1, 4). Most extra-synaptic GABA_A_ receptors contain α5 or δ subunits. These subunits are absent from postsynaptic GABA_A_ receptors, and their presence is thought to confer high affinity, which permits detection of micro-molar concentrations of ambient GABA, and low efficacy, which enables high potential for allosteric modulation(1, 4). A curious feature of the current passed by extra-synaptic GABA_A_ receptors is outward rectification at supra-threshold membrane voltage(5–11). Outward rectification enables greater inhibitory (hyperpolarising) current to flow during action potential (AP) generation, however its precise impact upon neuronal function is unclear. The extent of rectification may vary with GABA_A_ receptor subunit composition and is thought to be due to a voltage gating mechanism(12).

Tonic inhibition carries broad clinical and pharmacological significance(13). Extra-synaptic GABA_A_ receptors are a primary target of anaesthetic and anxiolytic agents, mutations cause inherited forms of epilepsy, and neurosteroid modulation can predispose to psychiatric disease or be utilised for therapeutic potential(4, 14). Recently, a positive allosteric modulator of extra-synaptic GABA_A_ receptors has shown benefit in post-partum depression and an antagonist is undergoing a phase 2 clinical trial in post-stroke motor recovery(15, 16).

Given the importance of tonic inhibition to neuronal function and its clinical relevance, there is much interest into how tonic inhibition modulates neuronal excitability(5, 6, 17–21). Neuronal excitability is often quantified using measures of rheobase and gain(17, 20, 21). Rheobase is defined as the minimum input that elicits an AP and gain the responsiveness of the neuron’s firing rate to changes of input. Within excitatory pyramidal cells, previous work has shown that tonic inhibition primarily modulates excitability by increasing rheobase(5, 17, 21). Non-rectifying inhibition, which obeys an ohmic (or linear) voltage-current relationship, can reduce gain during random stimuli that approximate *in-vivo* conditions by attenuating fluctuations in subthreshold input(17, 20, 21).

Within some inhibitory interneurons the presence of tonic inhibition has been observed to increase firing frequency(6, 7). This counterintuitive finding is explained by a depolarised GABA reversal potential relative to resting membrane potential, allowing extracellular GABA to raise membrane voltage closer to AP threshold and reduce rheobase(22). It is possible this excitatory influence of GABA may be restricted to certain subtypes of interneurons(6). This is of significance for understanding the impact of tonic inhibition upon network activity given the emerging role of interneuron subtypes for selectively fine tuning thalamo-cortical computations and neural information flow(23–25).

In light of this putative role of tonic inhibition in modulating interneuron function, the objective of this study was to characterise the impact of tonic inhibition upon the excitability of different subtypes of cortical interneurons. Interneuron subtypes may be classified according to expression of molecular markers (for example parvalbumin or somatostatin), their morphology, or their functional (electrophysiological) properties(26). With respect to their electrophysiological properties, interneurons may be broadly categorised into either fast spiking or non-fast spiking subtypes, although further fine-grained subcategories exist given by the Petilla nomenclature(27).

Using biophysically detailed computational models and patch-clamp recordings, we found that tonic inhibition reduced gain in most fast spiking interneurons, which are typically parvalbumin (Pv) positive, yet increased gain in most non-fast spiking interneurons that are typically somatostatin (Sst) positive. Differential gain modulation was observed between fast spiking and non-fast spiking interneurons. Analysis of our models revealed that differential gain modulation is enhanced by outward rectifying GABA_A_ receptors, occurs through a novel dendritic mechanism and is dependent upon intrinsic differences in neuronal AP dynamics.

## RESULTS

To explore the impact of inhibition upon interneuron excitability, we first generated biophysically-detailed interneuron models that were optimised to replicate the electrophysiological features of common subtypes (hereon referred to as ‘E-types’) of cortical interneurons (**Fig. 1a**). We generated ten models of a fast spiking interneuron (‘continuous non-accommodating’ or cNAC, using Petilla nomenclature) and ten models of two subtypes of non-fast spiking interneurons (‘continuous accommodating’ and ‘burst-accommodating’, or cAC and bAC respectively, **Fig. 1e**).

**Fig. 1:**
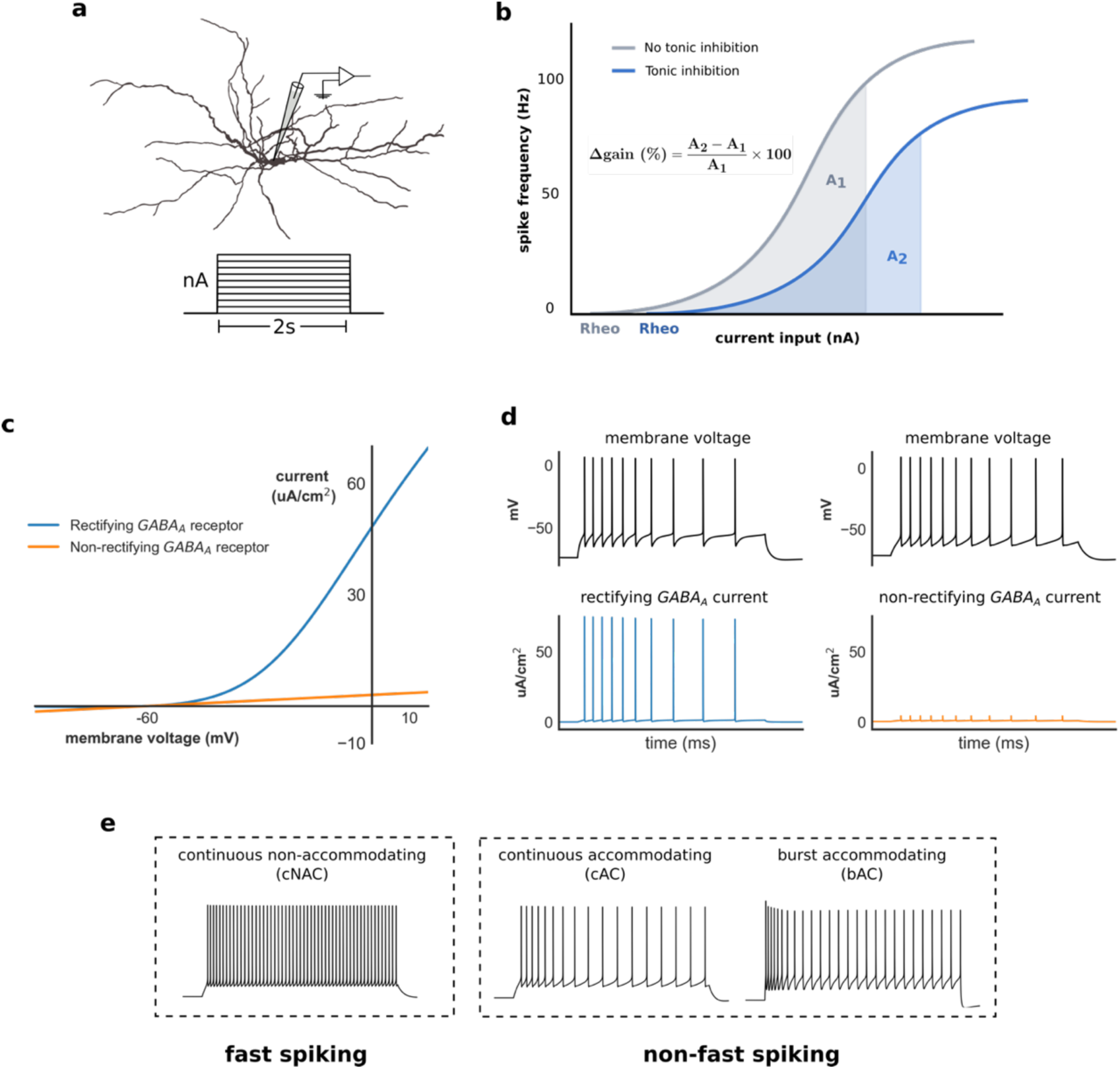
Computational modelling of interneuron excitability & tonic inhibition. **a)** Interneuron model morphology and step-current protocol used to obtain the current-frequency relationship. **b)** The current-frequency relationship was obtained with (blue) and without (grey) tonic inhibition, and the change in gain (Δ gain) defined as the relative difference in area under the current-frequency curve across a fixed input range (see Methods, *rheo* denotes rheobase). **c)** Voltage-current relationship of rectifying (blue) and non-rectifying (orange) extra-synaptic *GABA*_*A*_ receptors. **d)** Rectifying extra-synaptic *GABA*_*A*_ receptors allow greater outward (hyperpolarising) current to be passed at transmembrane voltages above ∼ −50mV, such as during AP generation. **e)** Time-voltage traces of three models optimised to exhibit different interneuron E-type classified according to either fast spiking vs non-fast spiking categories, or Petilla E-type. Note: by convention outward (hyperpolarising) transmembrane current is positive.

To determine the impact of tonic inhibition upon neuronal gain, the change in gain (Δ gain) in the presence of tonic inhibition was calculated for each model (**Fig. 1b**, see Methods). To examine the impact of outward rectification of extra-synaptic GABA_A_ receptors upon neuronal gain, Δ gain was calculated using both rectifying and non-rectifying tonic inhibition (**Fig. 1c & 1d**). Finally, since the reversal potential of GABA (*E*_*GABA*_) is known to influence interneuron excitability, Δ gain was also calculated for *E*_*GABA*_ of −60 and −80mV.

### 1) Tonic inhibition differentially modulates gain in functional subtypes of interneuron models

Consistent with previous studies exploring the influence of an inhibitory conductance upon neuron excitability(17), the presence of non-rectifying tonic inhibition generated an insignificant change in gain within non-fast spiking models for *E*_*GABA*_ of −60mV. A small reduction of Δ gain was observed in fast-spiking models (−6.0% ± 1.1, one-sample *t*-test, *t*(9) = −5.3, **P < 0.001, **Fig. 2a**) and there was a small but significant difference in Δ gain between fast spiking and non-fast spiking models (−6.0% ± 1.1 vs −0.8% ± 0.9, Welch’s *t*-test, *t*(28) = −3.4, *P* < 0.01). Similar findings were observed for *E*_*GABA*_ of −80mV, although reductions of Δ gain were observed in both fast spiking and non-fast spiking models (**Supplementary Fig. 1a**).

**Fig. 2:**
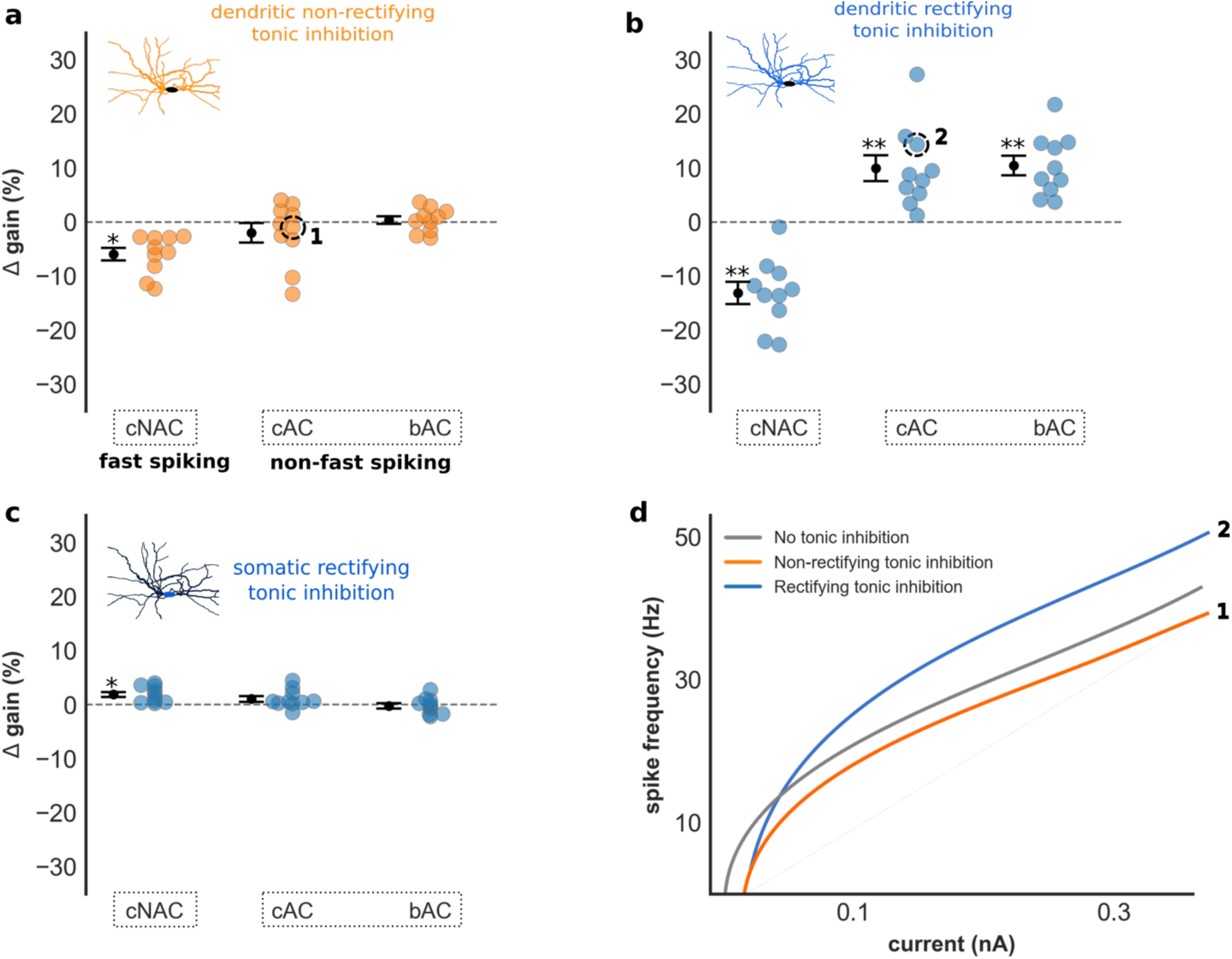
Impact of tonic inhibition upon gain in interneuron models. **a-c)** Δ gain of all interneuron models grouped by E-type (*E*_*GABA*_ = −60mV) for non-rectifying (**a**) and rectifying (**b**) tonic inhibition acting upon the dendritic tree, and rectifying tonic inhibition acting upon the soma (**c**). **a)** Non-rectifying tonic inhibition had insignificant impact upon gain in non-fast spiking models, and reduced gain in fast spiking models. A small but significant difference in Δ gain was observed between fast spiking and non-fast spiking models (−6.0% ± 1.1 vs −0.8% ± 0.9, Welch’s *t*-test, *t*(28) = −3.4, *P* < 0.01). **b)** Dendritic rectifying tonic inhibition increased gain in non-fast spiking models, and differentially modulated gain between fast spiking and non-fast spiking models (−13.1% ± 2.0 vs 10.2% ± 1.5, Welch’s *t*-test, *t*(28) = −9.32, *P* < 0.001). **c)** The presence of tonic inhibition at the soma did not induce differential gain modulation between E-types, suggesting gain modulation is mediated through a dendritic mechanism. **d)** Current-frequency relationship of a non-fast spiking model demonstrating changes in gain with non-rectifying (orange, labelled **1**) and rectifying (blue, labelled **2**) tonic inhibition. Results presented as mean ± s.e.m. Asterix denotes a significant Δ gain value compared to Δ gain = 0%, one-sample *t*-test, *\*P* < 0.01, *\*\*P* < 0.001.

Remarkably, we found that the presence of rectifying tonic inhibition increased gain within non-fast spiking models, and induced large differences in Δ gain between fast spiking and non-fast spiking models (−13.1% ± 2.0 vs 10.2% ± 1.5, Welch’s *t*-test, *t*(28) = −9.32, *P* < 0.001; **Fig. 2b**). A difference in Δ gain between fast spiking and both non-fast spiking Petilla E-types was also observed (one-way ANOVA with *post hoc* Tukey test, *F*(2,27) = 23.6, *P* < 0.001). Rectifying tonic inhibition, therefore, differentially modulated the gain of fast spiking and non-fast spiking interneuron models. This excitatory influence upon non-fast spiking models was also found for *E*_*GABA*_ of −80mV (**Supplementary Fig. 1b**). Interestingly an *E*_*GABA*_ of −80mV, which is hyperpolarised relative to the resting membrane potential of our models (**Supplementary Fig. 1f**), increased Δ gain in four models compared to *E*_*GABA*_ of −60mV. A hyperpolarised *E*_*GABA*_ relative to resting membrane potential is usually associated with reductions of neuronal excitability through increased rheobase, whereas our findings suggest a mixed inhibitory (increased rheobase) and excitatory (increased gain) influence.

Tonic inhibition is thought to be mediated by ambient GABA acting diffusely upon the spatial extent of the neuronal membrane(1, 28). Given our unexpected finding of differential gain modulation between fast spiking and non-fast spiking models, we next investigated how the spatial distribution of tonic inhibition impacts neuronal excitability. This was achieved by restricting tonic inhibition to different subcellular regions of each model, and then recalculating Δ gain (see Methods). Interestingly, differential gain modulation was only observed if rectifying tonic inhibition acted upon the dendritic tree (**Fig. 2b** & **Supplementary Fig. 1b & 1d**). In contrast, if tonic inhibition was restricted to the soma a significant difference in Δ gain between fast spiking and non-fast spiking models was not observed (1.9% ± 0.5 vs 0.4% ± 0.4, Welch’s *t*-test, *t*(28) = 2.4, n.s., **Fig. 2c** & **Supplementary Fig. 1c**) and increases of gain within non-fast spiking models did not occur. These findings suggest that rectifying tonic inhibition differentially modulates gain, and can enhance gain in non-fast spiking models, primarily by acting upon the dendritic tree.

Since neurons *in-vivo* are exposed to random synaptic input rather than constant current injections often used during experimental recordings, we next investigated the impact of tonic inhibition upon gain in response to noisy input conditions (**Supplementary Fig. 2a**, see Methods)(29). Again, rectifying tonic inhibition increased gain in non-fast spiking models and induced large differences in Δ gain between fast spiking and non-fast spiking models for *E*_*GABA*_ of −60mV (−9.3% ± 2.2 vs 6.1% ± 1.5, Welch’s *t*-test, *t*(28) = −5.7, \*\**P* < 0.001). A significance difference in Δ gain was observed between fast spiking and both non-fast spiking Petilla E-types (one-way ANOVA with *post hoc* Tukey test, *F*(2,27) = 18.5, \*\**P* < 0.001, **Supplementary Fig. 2b**). The presence of non-rectifying tonic inhibition also induced significant differences in Δ gain between fast spiking and non-fast spiking models, however differences were lower compared to rectifying tonic inhibition and increased gain within non-fast spiking models not observed (−5.4% ± 1.0 vs −0.7% ± 0.4, Welch’s *t*-test, *t*(28) = −4.4, \*\**P* < 0.001). We observed similar findings for *E*_*GABA*_ of −80mV (**Supplementary Fig. 2c**).

These findings reveal that tonic inhibition can differentially modulate gain within neuron models optimised to replicate features of different cortical interneuron E-types. Most surprisingly, tonic inhibition increased gain in non-fast spiking models. In our models, gain modulation is enhanced by outward rectifying *GABA*_*A*_ receptors, appears to occur through a dendritic mechanism and is dependent upon the model’s intrinsic electrophysiological properties.

### 2) Tonic inhibition differentially modulates gain in layer 2/3 cortical interneurons

In light of our modelling results, we next investigated if tonic inhibition can differentially modulate gain within interneuron subtypes of mouse somatosensory cortex. This was achieved by performing current-clamp recordings from layer 2/3 Sst-positive (n=11) and Pv-positive (n=10) interneurons. Each interneuron was then classified as fast spiking or non-fast spiking, and also according to its Petilla E-type.

We identified five distinct Petilla interneuron E-types from our recorded cells (**Fig. 3b**): cNAC (n=2), delayed non-accommodating (dNAC, n=6), cAC (n=9), non-adapting non-fast spiking (naNFS, n=3) and burst-irregular (bIR, n=1). cNAC and dNAC interneurons are considered fast spiking and all were Pv-positive. cAC and naNFS are considered non-fast spiking and, apart from one cAC interneuron, all were Sst-positive. Although bIR interneurons may be considered non-fast spiking, their precise classification is uncertain and so this interneuron was considered separately in our analysis(30).

**Fig. 3:**
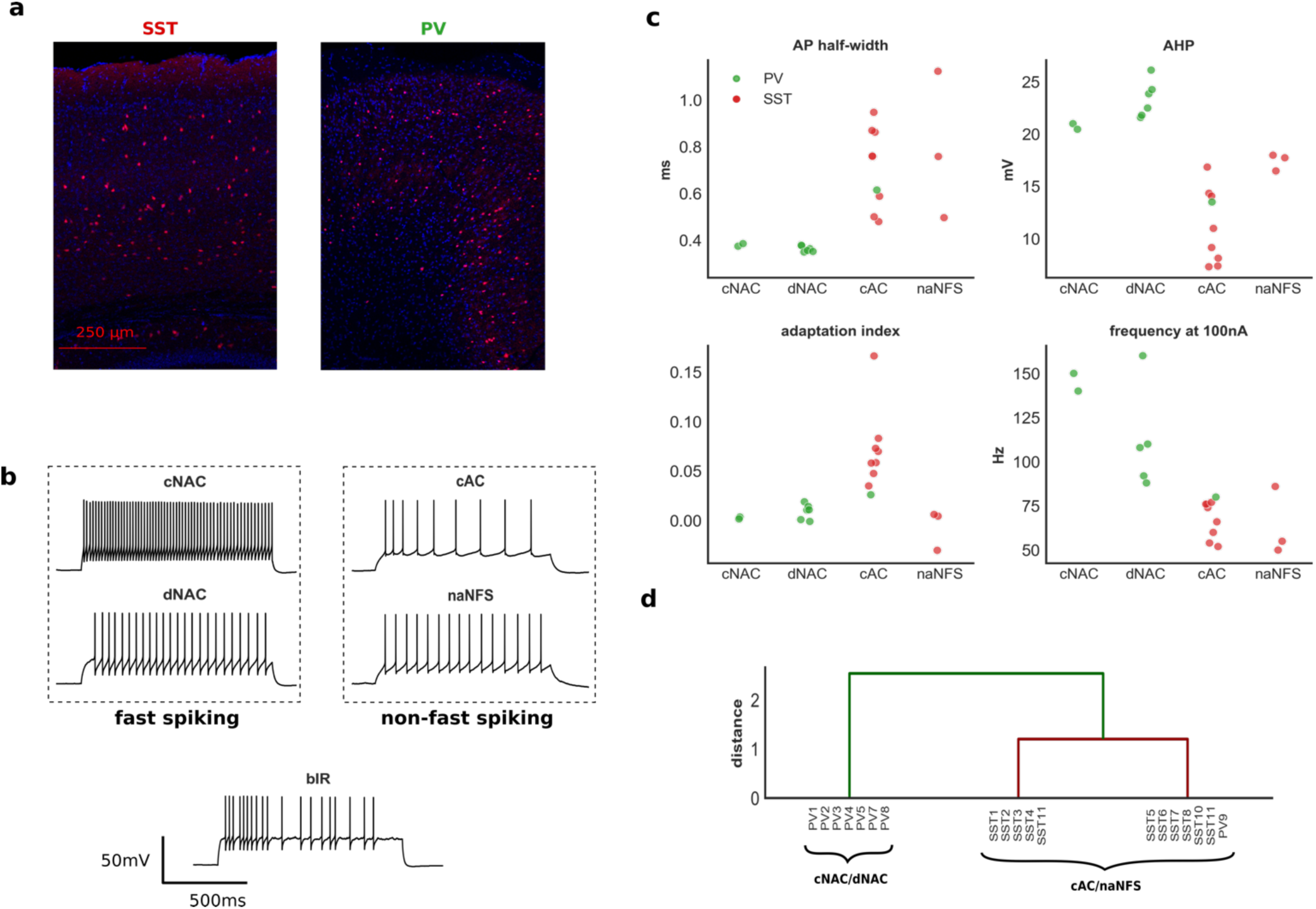
E-type classification of layer 2/3 cortical interneurons. **a)** Cortical immunohistochemical stain taken from an Sst (left) and Pv (right) positive mouse. **b)** Time-voltage traces from five recorded interneurons classified by E-type (all recordings shown in **Supplementary Fig. 3**). **c)** Four electrophysiologic features of recorded interneurons grouped by Petilla E-type. These features were used for hierarchical clustering (two Pv interneurons were excluded from this analysis, see Methods). **d)** Hierarchical clustering of recorded interneurons. Two major groups were identified: one consisting of cNAC & dNAC Petilla E-types, the other of cAC & naNFS E-types. This separation is supportive of our subjective Petilla classification and a division into fast spiking and non-fast spiking categories. AHP: afterhyperpolarisation; frequency at 100nA: spike frequency at 100nA above rheobase.

To test the validity of our classification, we performed hierarchical clustering on the recorded interneurons using four electrophysiologic features known to discriminate between fast spiking and non-fast spiking E-types (**Fig. 3c**, see Methods). The bIR and one dNAC interneuron were excluded from this analysis (see Methods). We identified greatest separation between two groups: one group containing all cNAC and dNAC E-types, the other containing cAC and naNFS E-types (**Fig. 3d**). This separation supports both our subjective Petilla classification, and our division of these interneurons into fast spiking and non-fast spiking respectively(30).

We then determined the impact of tonic inhibition upon neuronal gain by recording the current-frequency relationship of all interneurons before and after blockade of extra-synaptic GABA_A_ receptors (see Methods). Current-frequency curves were fitted using a Hill-type function and Δ gain calculated using a similar method to our models. In contrast to previous studies investigating the impact of tonic inhibition upon the gain of excitatory pyramidal cells, we observed a wide range of Δ gain values in recorded interneurons (**Fig. 4a**, time-voltage traces, current-frequency relationships and electrophysiologic characteristics of all cells in **Supplementary Fig. 3** & **Table 1**). Similar to our modelling results, tonic inhibition increased Δ gain within non-fast spiking interneurons (13.8% ± 3.6, one-sample *t*-test, t(7)=3.9, *P* < 0.01) and differentially modulated Δ gain between fast spiking and non-fast spiking interneurons (two-sided Mann-Whitney U test, *U = 0*, \*\*\**P* < 0.001, **Fig. 4a**). Importantly, we observed increased Δ gain a Pv-positive cAC interneuron, suggesting that increases of Δ gain are not dependent upon other neuronal properties determined by molecular marker(26). We also observed a large increase in Δ gain within the bIR interneuron.

**Table 1.**
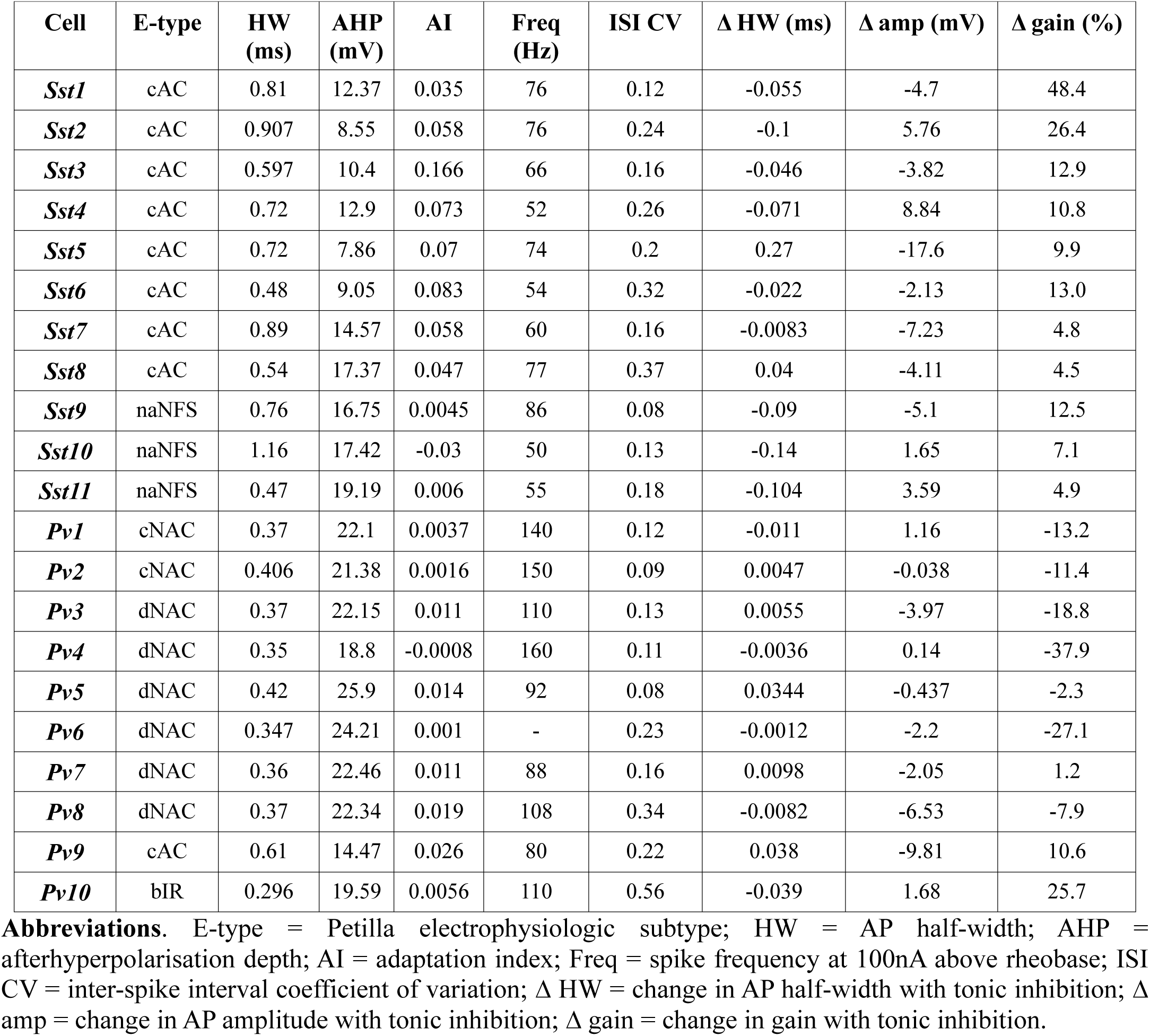
Electrophysiologic characteristics of recorded interneurons.

| Cell | E-type | HW (ms) | AHP (mV) | AI | Freq (Hz) | ISI CV | $\Delta$ HW (ms) | $\Delta$ amp (mV) | $\Delta$ gain (%) |
| --- | --- | --- | --- | --- | --- | --- | --- | --- | --- |
| *Sst1* | cAC | 0.81 | 12.37 | 0.035 | 76 | 0.12 | -0.055 | -4.7 | 48.4 |
| *Sst2* | cAC | 0.907 | 8.55 | 0.058 | 76 | 0.24 | -0.1 | 5.76 | 26.4 |
| *Sst3* | cAC | 0.597 | 10.4 | 0.166 | 66 | 0.16 | -0.046 | -3.82 | 12.9 |
| *Sst4* | cAC | 0.72 | 12.9 | 0.073 | 52 | 0.26 | -0.071 | 8.84 | 10.8 |
| *Sst5* | cAC | 0.72 | 7.86 | 0.07 | 74 | 0.2 | 0.27 | -17.6 | 9.9 |
| *Sst6* | cAC | 0.48 | 9.05 | 0.083 | 54 | 0.32 | -0.022 | -2.13 | 13.0 |
| *Sst7* | cAC | 0.89 | 14.57 | 0.058 | 60 | 0.16 | -0.0083 | -7.23 | 4.8 |
| *Sst8* | cAC | 0.54 | 17.37 | 0.047 | 77 | 0.37 | 0.04 | -4.11 | 4.5 |
| *Sst9* | naNFS | 0.76 | 16.75 | 0.0045 | 86 | 0.08 | -0.09 | -5.1 | 12.5 |
| *Sst10* | naNFS | 1.16 | 17.42 | -0.03 | 50 | 0.13 | -0.14 | 1.65 | 7.1 |
| *Sst11* | naNFS | 0.47 | 19.19 | 0.006 | 55 | 0.18 | -0.104 | 3.59 | 4.9 |
| *Pv1* | cNAC | 0.37 | 22.1 | 0.0037 | 140 | 0.12 | -0.011 | 1.16 | -13.2 |
| *Pv2* | cNAC | 0.406 | 21.38 | 0.0016 | 150 | 0.09 | 0.0047 | -0.038 | -11.4 |
| *Pv3* | dNAC | 0.37 | 22.15 | 0.011 | 110 | 0.13 | 0.0055 | -3.97 | -18.8 |
| *Pv4* | dNAC | 0.35 | 18.8 | -0.0008 | 160 | 0.11 | -0.0036 | 0.14 | -37.9 |
| *Pv5* | dNAC | 0.42 | 25.9 | 0.014 | 92 | 0.08 | 0.0344 | -0.437 | -2.3 |
| *Pv6* | dNAC | 0.347 | 24.21 | 0.001 | - | 0.23 | -0.0012 | -2.2 | -27.1 |
| *Pv7* | dNAC | 0.36 | 22.46 | 0.011 | 88 | 0.16 | 0.0098 | -2.05 | 1.2 |
| *Pv8* | dNAC | 0.37 | 22.34 | 0.019 | 108 | 0.34 | -0.0082 | -6.53 | -7.9 |
| *Pv9* | cAC | 0.61 | 14.47 | 0.026 | 80 | 0.22 | 0.038 | -9.81 | 10.6 |
| *Pv10* | bIR | 0.296 | 19.59 | 0.0056 | 110 | 0.56 | -0.039 | 1.68 | 25.7 |

**Abbreviations.** E-type = Petilla electrophysiologic subtype; HW = AP half-width; AHP = afterhyperpolarisation depth; AI = adaptation index; Freq = spike frequency at 100nA above rheobase; ISI CV = inter-spike interval coefficient of variation; $\Delta$ HW = change in AP half-width with tonic inhibition; $\Delta$ amp = change in AP amplitude with tonic inhibition; $\Delta$ gain = change in gain with tonic inhibition.

**Fig. 4:**
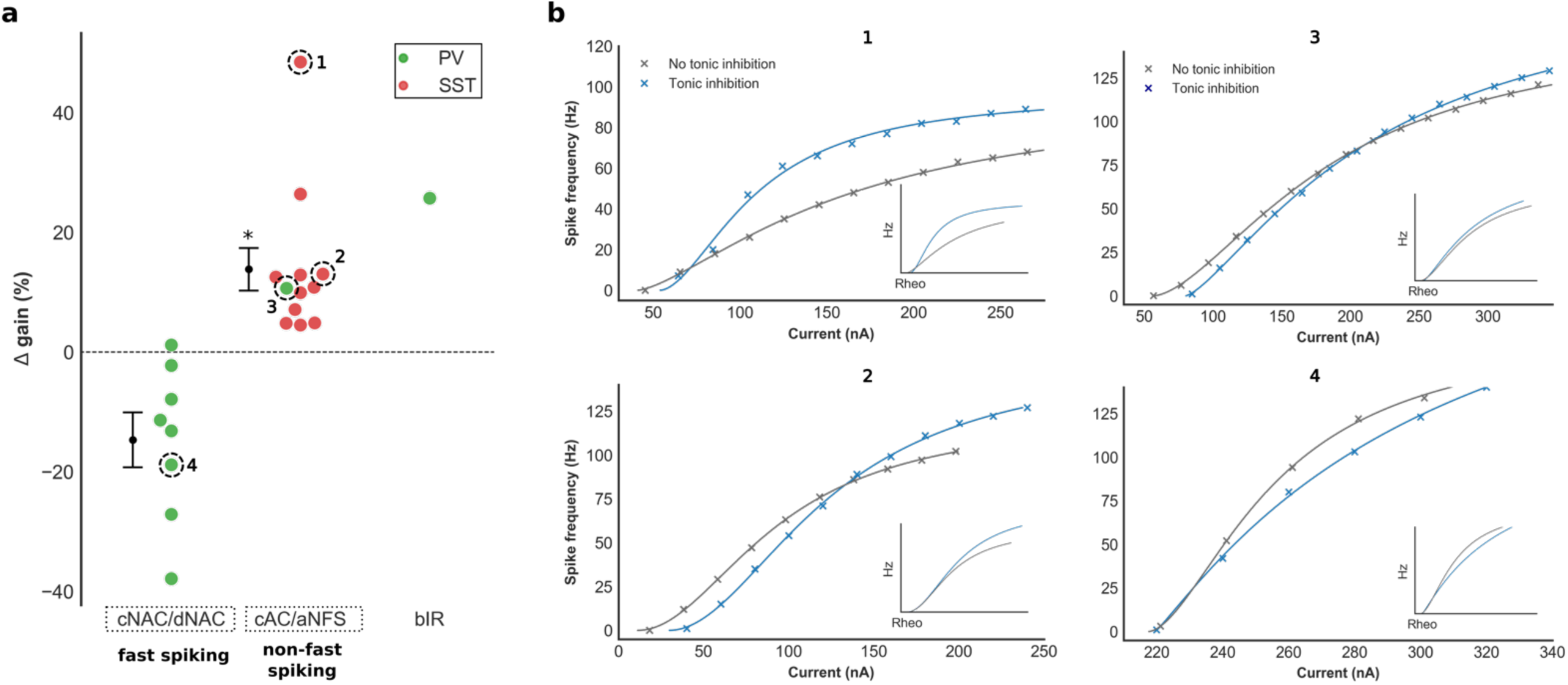
Impact of tonic inhibition upon gain in layer 2/3 cortical interneurons. **a)** Experimental changes in gain with tonic inhibition in fast-spiking (cNAC/dNAC), non-fast spiking (cAC/naNFS) and bIR interneuron, coloured by molecular marker (red: Sst, green: Pv). The current-frequency relationship of four interneurons are shown in **b** (inset: adjusted for rheobase). The presence of tonic inhibition increased gain in non-fast spiking compared to fast spiking interneurons (two-sided Mann-Whitney U test, *U = 0, P* < 0.001). Increased gain was observed in a non-fast spiking Pv interneuron (panel **3**) suggesting that differential gain modulation is not dependent upon other neuronal properties that vary with molecular marker. Tonic inhibition also increased gain within the bIR interneuron. Current-frequency relationships for all recorded interneurons are shown in **Supplementary Fig. 3**. Asterix denotes a significant Δ gain value compared to Δ gain = 0%, one-sample *t*-test, *\*P* < 0.01, *\*\*P* < 0.001.

Our models suggest that differential gain modulation is enhanced by the presence of outward rectifying GABA_A_ receptors. Since the changes in gain that we observed in our recordings were of a similar magnitude to those induced by rectifying tonic inhibition in our models (for example, compare Δ gain values in **Fig. 4a** with **Fig. 2b**), we obtained the voltage-current relationship of 9 Sst interneurons before and after blockade of extra-synaptic GABA_A_ receptors with picrotoxin (see Methods). Here, the subtracted voltage-current curve denotes the picrotoxin-sensitive component passed by extra-synaptic GABA_A_ receptors. We observed wide variation in the voltage-current relationship at depolarised membrane potentials (**Supplementary Fig. 4a**) and four interneurons displayed marked outward rectification.

### 3) Tonic inhibition enhances action potential repolarisation and reduces voltage-dependent potassium current at the soma

Our modelling and experimental results challenge the prevailing view that tonic inhibition has no impact upon neuronal gain during constant current stimuli and reduces gain during random stimuli(17). Surprisingly, we found that tonic inhibition can in fact *increase* gain within a potentially large subpopulation of inhibitory interneurons. To understand how tonic inhibition can exert this counterintuitive effect upon neuronal excitability, we recorded the total membrane current at the soma of our models during steady-state firing, with and without the presence of dendritic tonic inhibition (**Fig. 5a & 5b**). Total membrane current is equivalent to the sum of all ionic and axial currents that contribute to changes in somatic membrane voltage (**Fig. 5f** right, see Methods).

**Fig. 5:**
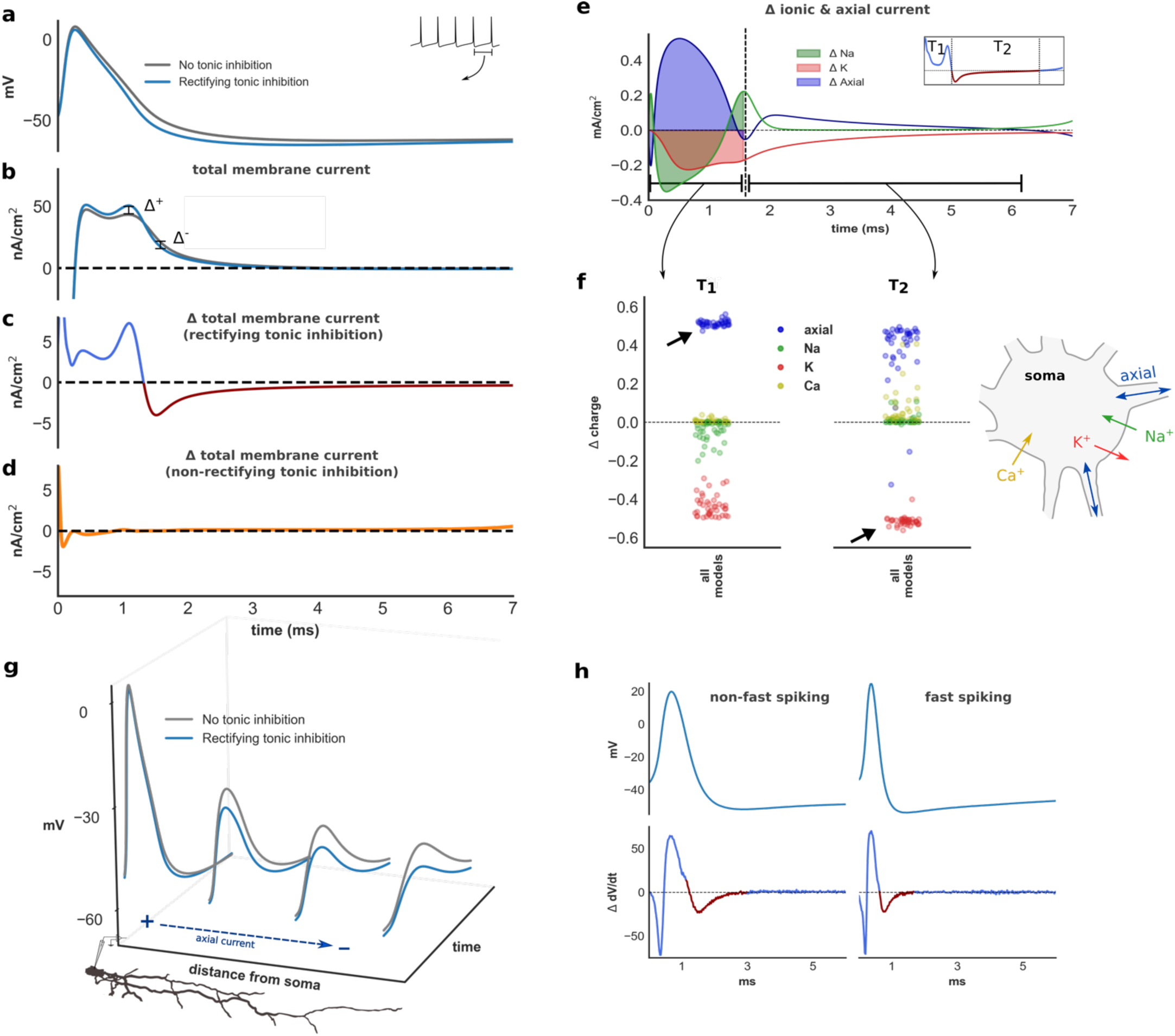
Biophysical impact of tonic inhibition. **(a)** Time-voltage trace during an inter-spike interval (ISI) in a model with (blue) and without (grey) dendritic rectifying tonic inhibition. Corresponding trace of total membrane current at the soma **(b)** and Δ total membrane current with rectifying **(c)** and non-rectifying **(d)** tonic inhibition. Rectifying tonic inhibition enhances AP repolarisation (**c**, blue trace T_1_ and corresponding to positive Δ) and promotes earlier recovery from repolarisation, evident by a depolarising change in total membrane current during AP downstroke (**c**, red trace T_2_ and corresponding to negative Δ). **e)** Changes in axial and ionic membrane current contributing to Δ total membrane current (K and Na refer to all potassium and sodium conductance’s. Leak and Ca current excluded since their contribution is minimal). **f)** Normalised change in charge (Δ charge) for each ionic species during period T_1_ and T_2_. Enhanced AP repolarisation is overwhelmingly mediated by increased somato-dendritic axial current. The biophysical basis for this is shown in **g**: rectifying tonic inhibition attenuates the electrotonic spread of an AP, increases somato-dendritic voltage gradient and enhances axial current flow during AP generation. In contrast, early recovery from AP repolarisation is overwhelmingly mediated by reductions of (hyperpolarising) potassium current (**f**). Furthermore, reductions of potassium current are observed throughout the ISI (see also **Supplementary Fig. 4**). **h)** Time-voltage trace and Δ total membrane current taken from a recorded fast spiking and non-fast spiking interneuron. Δ total membrane is estimated from the derivative of experimental time-voltage recordings (Δ dV/dt, see Methods).

We found that the presence of rectifying tonic inhibition increased the magnitude of outward (hyperpolarising) total membrane current during action potential (AP) generation (blue trace in **Fig. 5c**, all models shown in **Supplementary Fig. 4c**). In contrast, non-rectifying tonic inhibition produced minimal changes of Δ total membrane current during AP generation (**Fig. 5d**). Rectifying tonic inhibition, in other words, enhanced AP repolarisation. Consistent with this observation, rectifying tonic inhibition reduced either AP half-width (−0.037ms ± 0.005) or amplitude (−3.0mV ± 0.37) across all models (**Supplementary Fig. 1e**). To determine how rectifying tonic inhibition enhances AP repolarisation, we calculated the changes of current attributed to each ionic species at the soma (**Fig. 5e**). We then calculated changes of charge (Δ charge) deposited across the membrane for each species during enhanced AP repolarisation (period T_1_ in **Fig. 5e** & **5f**). We found that enhanced AP repolarisation was overwhelmingly mediated by increases of somato-dendritic (axial) current flow (**Fig. 5f**). The biophysical basis for increased somato-dendritic current is demonstrated in **Fig. 5g**. Here, the presence of rectifying tonic inhibition attenuates electrotonic spread of AP’s down the dendritic tree, as previously experimentally demonstrated in pyramidal neurons(28). This enhances the somato-dendritic voltage gradient and, by Ohm’s law, also increases somato-dendritic current flow.

In addition to enhancing AP repolarisation, we also observed that rectifying tonic inhibition promoted earlier recovery from AP repolarisation. This is reflected by an inward (depolarising) change in total membrane current during the AP downstroke and afterhyperpolarisation (AHP, outlined in red in **Fig. 5c** & **Supplementary Fig. 4c**). Using the same approach, we found that early recovery from AP repolarisation was overwhelmingly mediated by reductions of (hyperpolarising) potassium current (period T_2_, **Fig. 5f**). Notably, we also observed that tonic inhibition reduced voltage-dependent potassium current (and charge transfer) *throughout* the inter-spike interval (**Fig. 5f**, changes of all individual potassium currents for one model shown in **Supplementary Fig. 5b**). This is not surprising given the impact of rectifying tonic inhibition upon AP morphology: reductions of AP height and half-width afford less opportunity for the activation of voltage-dependent potassium channels. These findings are likely to confer a significant impact upon neuronal excitability. Alterations in transmembrane current flow during the AHP, largely due to changes in potassium channel activation, have previously been associated with changes of neuronal gain(31–33). Since reductions of potassium current are due to alterations of axial current flow secondary to attenuation of AP dendritic spread, this mechanism also provides an explanation for why gain modulation was only observed when tonic inhibition acted upon the dendritic tree.

To investigate if tonic inhibition produced a similar impact upon current flow at the soma in our experimental recordings, Δ total membrane current was estimated using the first derivative of the time-voltage trace (see Methods). Within most interneurons an increase in hyperpolarising current was observed during AP generation (**Fig. 5h & Supplementary Fig. 3**). Similar to our models, and consistent with our observation of enhanced AP repolarisation, reductions of either AP half-width or amplitude were observed in all recordings (**Table 1**). A depolarising change in total membrane current during AP downstroke and AHP was also frequently seen (**Fig. 5h & Supplementary Fig. 3**).

### 4) Tonic inhibition differentially modulates interneuron gain through variations in magnitude and deactivation kinetics of potassium current

We have demonstrated two biophysical consequences of rectifying tonic inhibition: enhanced AP repolarisation and reductions of transmembrane potassium current. These changes, however, were seen across all models. It remains unclear how these biophysical consequences produce gain modulation, and why differential gain modulation is observed between E-types.

To address this question, we created simplified single-compartment models of fast-spiking (cNAC) and non-fast spiking (bAC) interneurons (see Methods). The dynamics of each simple model are governed by three variables: fast (*v*, responsible for AP upstroke), slow (*w*, contributing to AP repolarisation and the inter-spike interval) and ultraslow (*u*, responsible for a voltage-dependent muscarinic potassium current (I_m_) that mediates spike-frequency adaptation). Each model was optimised to reproduce the electrophysiologic features and current-frequency relationship of a detailed model with and without rectifying tonic inhibition (**Fig. 6a**). The simple models with tonic inhibition also exhibited similar changes in total membrane current: enhanced AP repolarisation and early recovery from AP repolarisation (**Fig. 6a**). We found that features of different E-types could only be reproduced if different kinetics were used to describe the time course of the slow (*w*) variable. For the non-fast spiking model kinetics were based on activation of the persistent potassium (K_P_) current, and for the fast spiking model kinetics were based on activation of Kv_3.1_ (see Methods). We identified two mechanisms through which differential gain modulation can occur.

**Fig. 6:**
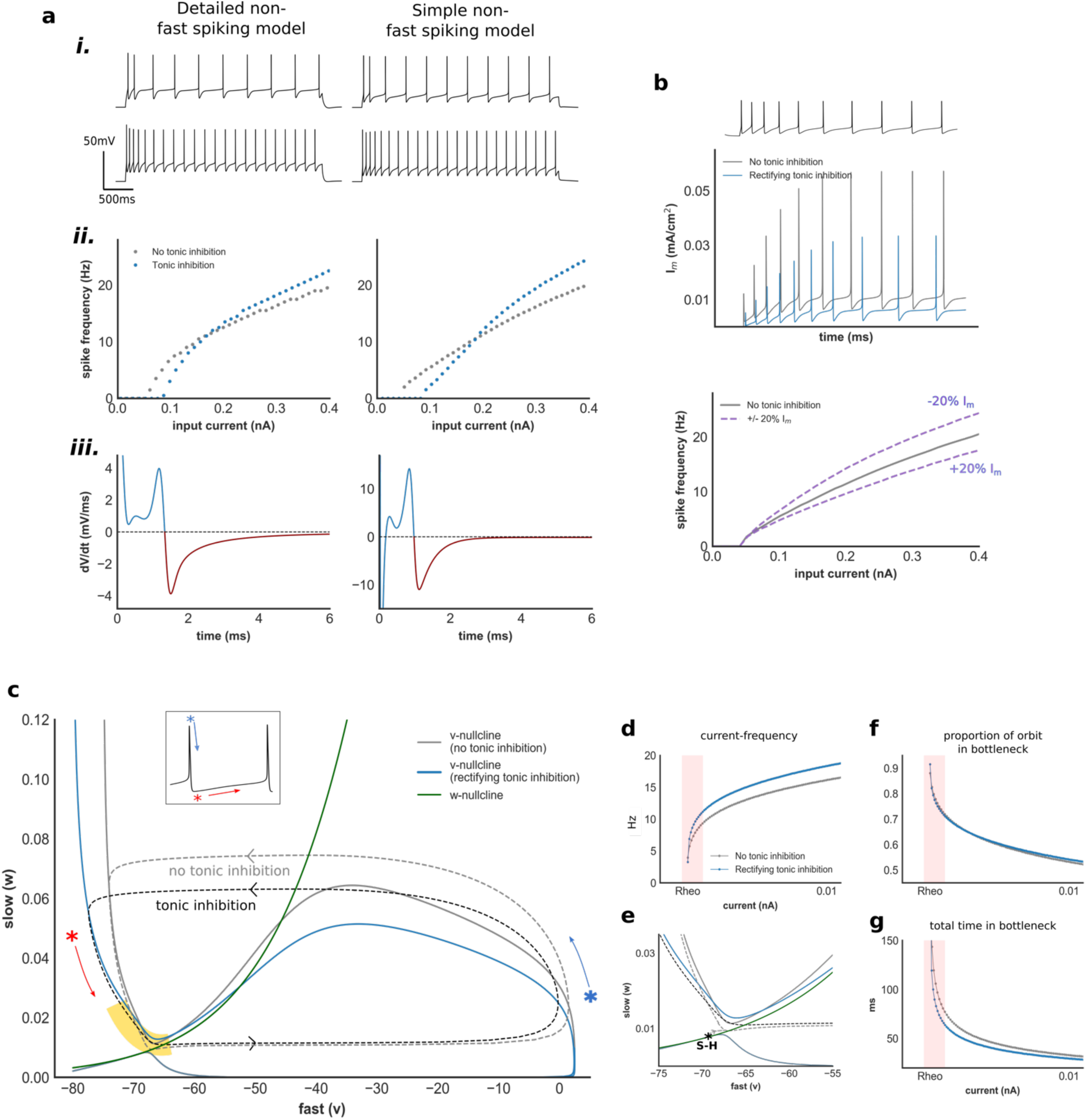
Impact of tonic inhibition upon AP dynamics within simplified non-fast spiking interneuron model. **a)** Electrophysiologic features (***i***), current-frequency relationship (***ii***) and Δ total membrane current (***iii***) within a simplified non-fast spiking model and its detailed counterpart (simplified fast spiking model in **Supplementary Fig. 5a**). **b)** I_m_ current generated by the non-fast spiking model without (grey) and with (blue) tonic inhibition, and impact of changes of I_m_ current upon gain in this model (c/w **Supplementary Fig. 5b**). **c)** Phase-plane and orbits during AP generation of the *v*-*w* subsystem of the non-fast spiking model, without and with tonic inhibition. Yellow denotes the bottleneck region. Blue/red asterix denote AP repolarisation and AHP, respectively, corresponding to inset time-voltage trace. **d)** Current-frequency relationship of the *v*-*w* system. Despite the absence of an ultraslow (I_m_) current, tonic inhibition increases gain. **e)** Phase-plane of the *v*-*w* system at the bottleneck. Transition from rest to spiking occurs via saddle-homoclinic (S-H) bifurcation (c/w **Supplementary Fig. 5e**). Tonic inhibition reduces activation of *w* and alters the trajectory through this region. Proportion of total orbit (**f**) and time (**g**) spent within bottleneck with increasing current input. Despite a similar proportion of the orbit spent within the bottleneck, the presence of tonic inhibition produced faster current-frequency scaling (pink region). This is reflected in a change in the eigenvalue of the unstable manifold of the S-H bifurcation (0.027 and 0.04 without and with tonic inhibition, respectively).

First, we found that tonic inhibition reduced I_m_ current in both fast spiking and non-fast spiking models due to reductions of AP height and AP half-width (**Fig. 6b & Supplementary Fig. 5b**). However, I_m_ current is roughly an order of magnitude greater in the non-fast spiking compared to fast spiking model (**Fig. 6b**). This is due to a higher channel density (*G*_*Im*_) required to generate spike-frequency adaptation in the non-fast spiking model. Crucially, varying the magnitude of the I_m_ current modulates gain (**Fig. 6b**) consistent with the known influence of a slow adapting current upon the neuronal input-frequency relationship(34). Here, due to differences in magnitude of *G*_*Im*_ related to neuronal E-type, a similar proportional reduction of I_m_ current produces far greater increase in gain within the non-fast spiking model (**Supplementary Fig. 5d**).

Next, we examined the input-frequency relationship of the fast-slow (*v*-*w*) subsystem of each simplified model (**Fig. 6c & Supplementary Fig. 5**, see Methods). Despite the absence of an I_m_ current, we still observed differential gain modulation between E-types, suggesting that another mechanism is also contributing (**Fig. 6d**). Analysing the phase portraits of both models, we found that tonic inhibition reduces the activation of *w* which can be seen in a narrowing of the height of the orbit during AP generation (**Fig. 6c** & **Supplementary Fig. 5c**). Since *w* mediates AP repolarisation and the AHP, this corresponds to a reduction of voltage-dependent potassium current responsible for AP repolarisation in our detailed models. Within both E-types, the input-frequency relationship is initially dominated by the presence of a bottleneck near the bifurcation from rest to spiking that allows for low-frequency firing (**Fig. 6f** & **Supplementary Fig. 5f**, highlighted in yellow in **Fig. 6c** & **Supplementary Fig. 5c**). In the non-fast spiking model, *w* continues to deactivate slowly as the orbit traverses this bottleneck. Therefore, in the presence of tonic inhibition, reduced activation of *w* during AP generation alters the trajectory through this bottleneck (**Fig. 6e**). This enhances frequency scaling and increases gain (**Fig. 6g**).

In contrast, *w* deactivates rapidly during the AP downstroke and AHP within the fast spiking model. Rapid deactivation is due to faster deactivation kinetics associated with Kv_3.1_ channels that are thought to enable high frequency firing(26). Consequently, the orbit traverses the bottleneck adjacent to the *w* nullcline both with and without tonic inhibition (**Supplementary Fig. 5c** & **Supplementary Fig. 5e**). Tonic inhibition also reduces the activation of *w* during AP generation in the fast spiking model. However, this exerts minimal impact upon frequency scaling since the trajectory through the bottleneck remains unchanged (**Supplementary Fig. 5f** & **Supplementary Fig. 5g**).

To summarise, enhanced AP repolarisation and reductions of voltage-dependent potassium current differentially modulate gain via two distinct mechanisms. First, reductions of ‘ultraslow’ potassium current preferentially increase gain in non-fast spiking interneurons due to higher channel densities required to generate spike-frequency adaptation. Second, reductions of potassium current mediating AP repolarisation and the AHP modulate gain according to channel deactivation kinetics. If there is rapid channel deactivation through Kv_3.1_ – a potassium channel known to be strongly expressed within fast spiking interneurons(26) - reductions in potassium current exert minimal influence upon gain.

### 5) Changes of interneuron gain modulate gamma-frequency oscillations in a network model

Our modelling and experimental results have shown that tonic inhibition can unexpectedly modulate the gain of inhibitory interneurons. Interestingly, interneuron gain modulation has also been shown to occur in response to other neuromodulators such as acetylcholine and serotonin(31, 32). Although gain modulation of excitatory pyramidal cells is thought to subserve a number of behaviourally-relevant changes in network activity, the influence of changes of interneuron gain upon activity within a neuronal network is less well established(35).

As a preliminary exploration of this question in light of our results, we explored the impact of interneuron gain within a network model consisting of excitatory (PC) and inhibitory (SST) neurons providing feedback inhibition (**Fig. 7a**)(36). At sufficient input amplitude (I_stim_) and recurrent excitatory conductance (*G*_*pp*_) this network exhibits transient and sustained gamma-frequency oscillations (**Fig. 7b**). We performed a sensitivity analysis to investigate the impact of 20% and 40% increases in SST gain. We found that increased SST gain produced a wider parameter range over which gamma oscillations could be sustained (**Fig. 7c**). Interestingly, it has recently been shown that activation of Sst interneurons can promote cortical gamma-frequency local field potentials(37).

**Fig. 7:**
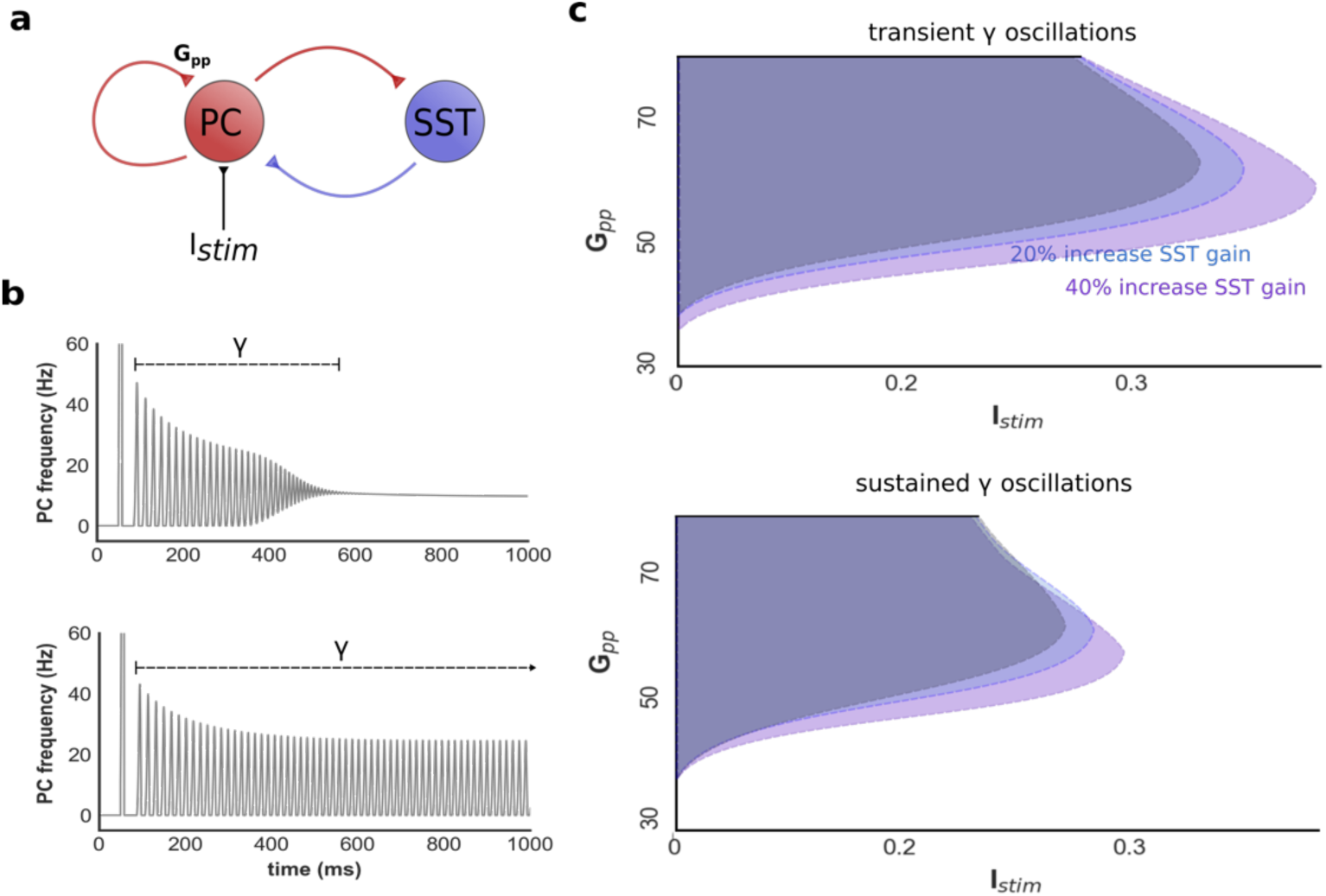
Impact of changes of interneuron gain upon gamma-frequency oscillations within a network model. **a)** Rate-based network consisting of pyramidal neurons (PC) with recurrent excitatory synapses (synaptic strength = *G*_*pp*_) and reciprocally connected interneurons (SST) providing feedback inhibition. **b)** With sufficient input (*I*_*stim*_) and recurrent excitatory synaptic strength this network generates transient (top) and sustained (bottom) gamma (γ)-frequency oscillations. **c)** Sensitivity analysis demonstrating range of *I*_*stim*_ and *G*_*pp*_ values over which transient (top) and sustained (bottom) gamma-frequency oscillations occur. Increased SST gain promotes network oscillations over a wider parameter range (grey: baseline, blue: 20% increase, purple: 40% increase).

## DISCUSSION

Using biophysically-detailed neuron models and patch-clamp recordings we have shown that GABA-mediated tonic inhibition differentially modulates gain in cortical interneurons. Surprisingly, we found that tonic inhibition increases gain within interneurons possessing electrophysiological characteristics typical for Sst interneurons. Therefore, our results challenge the prevailing view that tonic inhibition either has no impact on, or reduces, neuronal gain(17, 19–21).

Remarkably, our results suggest that differential gain modulation is dependent upon the intrinsic electrophysiological properties of interneurons. This finding is of particular significance when considering the neuromodulatory action of ambient GABA. Ambient GABA diffuses throughout the extracellular space and fluctuations in concentration exert profound influence over the excitability of cortical neurons(2). These fluctuations are driven by GABA transporter activity, synaptic spill-over and volume transmission from neurogliaform cells(1, 38). Instead of providing a simple ‘blanket’ of inhibition, we reveal that non-specific diffusion of GABA can specifically tune interneuron excitability. Although other neuromodulators, such as acetylcholine, can mediate selective interneuron excitation via cell-to-cell variation in receptor expression, GABA can achieve a similar feat through cell-to-cell variation in electrophysiological properties(39, 40).

Given the crucial influence of GABA upon brain function, our results imply that differential excitation of interneuron E-types may promote behaviourally relevant network activity. A growing literature suggests that many neuromodulators regulate interneuron excitability to produce transitions in network activity(26, 40, 41). Selective modulation of interneuron subtypes, however, is usually defined according to the interneuron’s molecular marker, for instance through optogenetic manipulation of Pv or Sst-expressing interneurons(23, 25). Our results extend this concept to suggest that interneuron subtypes, instead defined by their electrophysiologic characteristics, can also undergo selective modulation *in-vivo*.

This study demonstrates that rectifying tonic inhibition exerts markedly different biophysical effects to non-rectifying inhibition. Non-rectifying tonic inhibition has previously been shown to have minimal impact upon neuronal gain(17, 19). Existing models suggest that during spiking activity, a non-rectifying conductance has little influence upon AP dynamics because it generates a small inhibitory current relative to the large sodium and potassium currents responsible for AP upstroke and repolarisation(17, 19). Our findings modify this picture in three ways. First, the presence of outward rectification creates a large hyperpolarising current during AP generation. Second, using neuron models with realistic dendritic morphology, we show that rectifying tonic inhibition attenuates the amplitude of the electrotonic spread of AP’s. Dendritic AP attenuation has previously been demonstrated within pyramidal neurons but, crucially, we show that it exerts strong influence upon AP dynamics at the soma through enhanced somato-dendritic ‘axial’ current(28). Notably, this hyperpolarising axial current reduces the activation of voltage-dependent potassium channels at the soma. Finally, by generating models with a range of electrophysiological properties we show that these biophysical effects can exert differential impact upon interneuron excitability.

We identified two properties that differ between fast spiking and non-fast spiking interneurons to enable tonic inhibition to differentially modulate gain. The first is the magnitude of an ‘ultraslow’ conductance that generates spike-frequency adaptation within non-fast spiking interneurons. Previous theoretic work has shown that an ultraslow, adapting current can reduce gain(34). Here, we show that tonic inhibition can deactivate such currents through changes in AP morphology to increase gain. The second relates to differences in deactivation kinetics of the repolarising potassium current. Our simplified fast spiking model could only reproduce features of its detailed counterpart if kinetics of the slow variable (*w*) were based upon Kv_3.1_. Notably, Kv_3.1_ is strongly expressed within fast spiking interneurons and deactivates rapidly during AP downstroke to enable high-frequency firing(42). Although tonic inhibition reduces the activation of *w*, this exerts minimal influence upon gain since *w de*activates rapidly during the AHP (**Supplementary Fig. 5c & 5e**). In contrast, *w* deactivates slowly in the non-fast spiking model. Therefore, reduced activation of *w* with tonic inhibition alters the trajectory through the bottleneck that dominates the current-frequency relationship. Such bottlenecks (or attractor ruins) are well described within neuron models that sustain low-frequency firing and occur following bifurcation from rest to spiking(43). Their initial input-frequency scaling (ie gain) is related to the eigenvalue associated with the unstable manifold at the bifurcation(43, 44). In the non-fast spiking model, slow *w* deactivation allows tonic inhibition to increase the eigenvalue of the unstable manifold and enhance gain. It is possible that these characteristics could be exploited through other pharmacologic means to enable gain modulation. Interestingly, acetylcholine and serotonin have also been shown to enhance gain and modify the AHP within non-fast spiking interneurons(31, 32).

Our results shed light on the functional relevance of outward rectifying extra-synaptic GABA_A_ receptors. Outward rectification has been repeatedly observed in experimental studies yet its significance to neuronal function unclear(5–11). Rectification may be dependent upon GABA_A_ receptor subunit composition and, in our study, we observed strong rectification in 4/9 recorded cells(45). This raises the possibility that differential gain modulation observed in this study could be further ‘enriched’ by targeting specific GABA_A_ receptor subtypes, for instance through neurosteroids(13). Furthermore, although we have investigated neocortical interneurons, it is possible that other brain regions such as the hippocampus may preferentially express rectifying GABA_A_ receptors(5).

It is probable that our results generalise across species and age. Our models were based on recordings from juvenile rat neocortex, yet our experimental results obtained from adult mice. We observed expected age-related electrophysiological differences between model and experiment(46). AP half-width, for instance, was significantly shorter in our slice recordings (compare AP morphology between **Fig. 5a** and **Fig. 5h**). Nevertheless, our model predictions held. This suggests that tonic inhibition may exploit conserved features of interneuron E-types to induce differential gain modulation.

Our findings raise a number of further questions. Tonic inhibition exerts strong influence upon neuronal excitability within the thalamus and a differential role for Sst and Pv interneurons in this region is beginning to emerge(23). Our findings raise the possibility that ambient GABA may promote thalamo-cortical activity preferentially driven by Sst-positive thalamic reticular nucleus interneurons. Interneurons that express the ionotropic 5HT3a receptor may exhibit similar electrophysiologic properties to Sst neurons but were not investigated in this study(26). We would anticipate similar results within this interneuron population. Since the biophysical effects of tonic inhibition were mediated through the dendritic compartment, we would also anticipate neuronal morphology to modify tonic inhibition-related gain modulation. We would therefore predict more extensive arborisation to enhance gain. Finally, the impact of interneuron gain modulation upon cortical network activity remains unclear. A simple rate-based model suggests a role in modulating oscillatory activity, but further studies are required to elucidate these network-level effects.

## Author Contributions

A.B., S.P., S.H., C.A.R., D.B.G and S.B. conceived the study and designed the experiments. A.B., B.Z. and C.R. performed detailed modeling. R.H. performed patch-clamp experiments. A.B. and R.H. analysed experimental data. A.B. and B.Z. performed simple and network modeling. A.B., S.P., C.A.R., S.H., D.B.G. and B.Z. wrote the manuscript.

## Acknowledgements

This work was supported, in part, through funding for the Blue Brain Project (BBP) from the ETH Domain.

## METHODS

### 1) Detailed neuron modelling & optimisation

A model of a layer 2/3 Basket Cell (L23BC) available through the Blue Brain Project Neocortical Microcircuit portal was used for single neuron optimisation and simulation(47). The model contains >200 compartments, a calcium diffusion mechanism and 11 voltage-dependent channel mechanisms based on a Hodgkin-Huxley formulation: fast (transient) sodium and potassium (Na_T_ & K_T_), persistent sodium and potassium (Na_P_ & K_P_), Kv_3.1_, M-current (I_m_), hyperpolarisation-activated current (I_h_), calcium-activated potassium (SK), high and low voltage-activated calcium (Ca_H_, Ca_L_) and a leak current (pas). Parameter values for each mechanism are found in *Markram et al* and references therein(48). Simulations and analysis were performed using NEURON and Python(49).

Rectifying tonic inhibition was modelled as a voltage-dependent conductance with channel kinetics based on *Pavlov et al*(5):

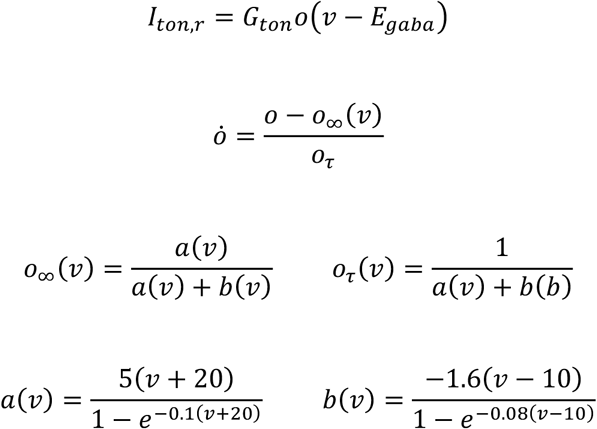

where *I*_*ton,r*_ is rectifying tonic current, *G*_*ton*_ peak conductance, *o* activation variable, *v* membrane voltage, *E*_*GABA*_ GABA reversal potential, *o*_*∞*_*(v)* and *o*_*τ*_*(v)* channel activation steady-state and time constant functions, respectively. Tonic inhibition was present in all compartments and *G*_*ton*_ retained as a free parameter during model optimisation(50).

*G*_*ton*_ upper and lower bounds were based on experimental changes in whole-cell holding current after extra-synaptic GABA_A_ receptor blockade(3). This was performed as follows. First, rectifying tonic inhibition was added to a previously optimised L23BC model. *E*_*GABA*_ was set to the chloride reversal potential used in the experimental setup. An *in-silico* voltage-clamp was applied at the soma and *G*_*ton*_ increased until ‘blockade’ (setting *G*_*ton*_ to 0 S/cm^2^) generated similar change in holding current to experimental results. The upper range of experimental results using this approach was 5×10^−4^S/cm^2^. Therefore, *G*_*ton*_ was restricted between 0 and 5×10^−4^ S/cm^2^ during optimization.

The peak conductance of ion channel mechanisms was optimised using a feature-based multi-objective algorithm implemented through BluePyOpt(51). Full details of this approach are outlined elsewhere(52, 53). Briefly, models were stimulated with somatic current and values of 32 to 42 electrophysiologic features extracted(54). For each feature, a Z-score was calculated based on *in-vitro* slice recordings from juvenile rat neocortex. These Z-scores were used as the fitness function for an evolutionary algorithm(52). Features were optimised to one of three Petilla electrophysiologic subtypes (E-types) of cortical interneurons: continuous accommodating (cAC), continuous non-accommodating (cNAC) and burst accommodating (bAC)(55, 56). Models were accepted if the sum of fitness scores was under 50. The optimisation algorithm was implemented on a computing cluster based at the Swiss National Supercomputing Centre. 10 models of each E-type (30 models total) were created. For each model, a different random seed was used to implement the optimisation algorithm.

### 2) Model stimulation & gain calculation

Input-frequency relationships of optimised models were obtained using constant current injected at the soma and a point process that generated excitatory conductance noise. Excitatory noise was modelled as an Ornstein-Uhlenbeck stochastic process with parameters based on *Destexhe et al*(29). Input-frequency curves were derived by increasing mean conductance and variance. To investigate the impact of rectifying tonic inhibition upon neuronal gain, input-frequency curves were calculated for two *G*_*ton*_ values: 0 and 0.001 S/cm^2^ (referred as ‘without’ and ‘with’ tonic inhibition hereon).

Change in gain (Δ gain) was calculated as the relative change in area under the input-spike frequency curve (AUC) over a fixed input range without and with tonic inhibition:

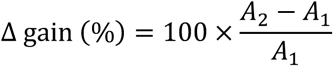

Where *A*_*2*_ denotes AUC with tonic inhibition, and *A*_*1*_ AUC without tonic inhibition (**Fig. 1b**). Δ gain was calculated using AUC rather than gradient because this provides a more consistent measure and allows for comparison between input-spike frequency curves despite variations in the shape of the input-spike frequency relationship.

The input range used to calculate AUC was defined from rheobase to an amplitude that elicited either spike-frequency of 100Hz or produced depolarisation block. The smallest range meeting these criteria with or without tonic inhibition was used. 100Hz was chosen as the upper limit because it encompasses a range of spike frequencies observed *in-vivo* for Pv and Sst-positive interneurons in awake animals(57, 58). Δ gain was calculated for *E*_*GABA*_ of both −80 and −60mV, and as *G*_*ton*_ was varied within different subcellular regions: the soma, dendritic tree and all compartments. Δ gain was also calculated after replacing rectifying tonic inhibition with a non-rectifying conductance:

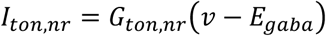

*I*_*ton,nr*_ denotes non-rectifying current and *G*_*ton,nr*_ non-rectifying peak conductance. *G*_*ton,nr*_ was set to a value that produced an identical change in holding current to *G*_*ton*_ after *in-silico* voltage clamp ‘block’ using an identical approach to Section 1.

### 3) Experimental animals

All experimental procedures in this study were conducted in accordance with the Prevention of Cruelty to Animals Act 1986, under the guidelines of the NHMRC Code of Practice for the Care and Use of Animals for Experimental Purposes in Australia and were approved by the Florey Neuroscience Institute Animals Ethics Committee. Homozygous Sst^tm2.1(cre)Zjh^/J mice (Jax Stock number: 013044) or B6;129P2-Pvalb^tm1(cre)Arbr^/J mice (Jax Stock 008069) were crossed with homozygous B6.Cg-Gt(ROSA)26Sor^tm14(CAG-tdTomato)Hze^/J mice (Jax Stock number: 007914) to produce mice that expressed tdTomato specifically in SST or PV expressing interneurons, hereafter termed SST-positive or PV-positive mice.

### 4) Brain slice preparation

Sst-positive and Pv-positive mice (6-8 weeks old, n = 3 mice) were anesthetized with 2% isoflurane and decapitated. The brain was removed quickly and placed into an iced slurry of cutting solution consisting of (mM): 125 Choline-Cl, 2.5 KCl, 0.4 CaCl2, 6 MgCl2, 1.25 NaH2PO4, 26 NaHCO3, 20 D-glucose saturated with 95% O2 plus 5% CO2. 300 μm coronal cortical slices were cut on a vibratome (VT1200; Leica; Germany) for whole-cell patch-clamp experiments. Slices were incubated at room temperature for a minimum of 1 hour in artificial cerebral spinal fluid (aCSF) consisting of (mM): 125 NaCl, 2.5 KCl, 2 CaCl2, 2 MgCl2, 1.25 NaH2PO4, 26 NaHCO3, 10 D-glucose, saturated with 95% O2 plus 5% CO2 before patching.

### 5) Whole-cell patch-clamp electrophysiology

Slices cut from Sst-positive and Pv-positive mice were transferred to a submerged recording chamber on an upright microscope (Slicescope Pro 1000; Scientifica, UK) and perfused (2 ml/min) with aCSF at 32 °C. Layer 2/3 interneurons were visually identified using fluorescence targeted patching with infrared-oblique illumination microscopy with a 40x water-immersion objective (Olympus, Japan) and a CCD camera (IEEE 1394; Foculus, Germany). Whole-cell patch-clamp recordings were made in voltage and current clamp modes using a PatchStar micromanipulator (Scientifica, UK) and an Axon Multiclamp 700B patch-clamp amplifier (MDS, USA). Data were acquired using pClamp software (v10; MDS, USA) with a sampling rate of 50 kHz and low pass Bessel filtered at 10 kHz (Digidata 1440a; Axon). Patch pipettes (4-7 MΩ; GC150F-10; Harvard Instruments, USA) pulled using a Flaming/brown micropipette puller (Model P-1000; Sutter Instruments, USA). For current clamp recordings, patch pipettes were filled with a solution consisting of (mM): 125 K-gluconate, 4 KCl, 2 MgCl_2_, 10 HEPES, 10 EGTA, and 0.3 GTP-Na (pH 7.3 and 300 mOsm). For voltage clamp recordings, CsCl (15 mM; Sigma, Australia), TEA (10mM; Torcris, Australia) and Qx314 (5 mM; Torcris, Australia) were added to the above internal solution. Extracellular blockade of α-amino-3-hydroxy-5-methyl-4-isoxazole-propionic acid (AMPA) and kinate receptors was achieved with 2,3-dihydroxy-6-nitro-7-sulfamoyl-benzo[f]quinoxaline-2,3-dione (NBQX; 50 µM; Sigma, Australia), N-methyl-D-aspartate (NMDA) receptors with (2R)-amino-5-phosphonopentanoate (AP5; 50 µM; Torcris, Australia), γ-aminobutyric acid B (GABA_B_) receptors with CGP5243 (25 µM; Sigma, Australia), phasic GABA_A_ receptors with SR95531 (0.5 µM; Sigma, Australia), and extrasynaptic (tonic) GABA_A_ receptors with picrotoxin (100 µM; Sigma, Australia). GABA (5 µM; Torcris, Australia) was included in the aCSF solution to provide a basal level of GABA agonism.

Once whole-cell configuration was obtained, to characterize firing a holding current was injected to maintain a membrane potential of approximately −70 mV and current steps were applied (−60 to 320 pA steps amplitude in 20 pA increments, 1 s step duration) in current clamp mode. To determine I-V relationships, cells were held at −70 mV and a voltage ramp (from −70 mV to −20 mV, 5 s duration) and voltage step (from −70 mV to 30 mV, 1 s duration) applied. Series resistance and whole-cell capacitance compensation were applied. To be included in the present study a cell had to have an access resistance of less than 20 MΩ and a holding current of less than −200 pA throughout the entire recording. Current and voltage clamp protocols were performed before and after application of picrotoxin. Voltage-dependence of the picrotoxin-sensitive current (ie extra-synaptic GABA_A_-mediated current) was calculated by subtracting the I-V relationship before and after picrotoxin. Values presented are not corrected for liquid junction potential. The I-V relationship in **Figure S3** is normalised for each cell relative to the picrotoxin-sensitive current at −70mV.

### 6) Experimental data analysis & E-type classification

Experimental data analysis was performed using Axograph X software (Berkeley, USA) and the Electrophys Feature Extraction Library (EFEL)(59). AP onset was defined when the first derivative of the voltage trace exceeded 12V/s for at least 5 consecutive time points. After-hyperpolarization potential (AHP) amplitude was calculated relative to voltage at AP onset. AP half-width (AP-HW) was measured at 50% of AP amplitude. Adaptation index (AI) was calculated as the normalised average difference of two consecutive inter-spike intervals (ISIs). Frequency at 100nA is defined as spike frequency at input current 100nA above rheobase. AP amplitude was calculated relative to minimum AHP voltage. The input resistance was calculated from the voltage deflection relative to baseline that occurred from injection of a −5 pA, 50 ms duration test pulse. Resting membrane potential was determined as the membrane voltage of a cell without injection of a holding current.

Recorded neurons were classified using the Petilla classification by three co-authors of this study with complete inter-observer agreement (AB, CR and SP). Electrophysiologic features of each neuron were extracted at rheobase (with the exception of frequency at 100nA) using the EfEL feature extraction library. Hierarchical clustering was performed using Ward’s method with four features: AP-HW, AHP, AI and frequency at 100nA. PV10 was excluded from analysis because it had clearly distinct features to eye compared to other recorded cells, and PV6 was excluded because a spike frequency 100nA above rheobase was not obtained with the experimental protocol.

The current-frequency relationship of each neuron was fitted using a Hill-type function(17):

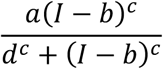

where *I* is input current and *a, b, c* and *d* free parameters. Δ gain was calculated using a similar approach to Section 2. To avoided misleading Δ gain values generated by the fitting procedure, the lower input range used to calculate AUC was defined as the minimum input that elicited a spike frequency above 1Hz.

### 7) Analysis of detailed models

Detailed models were analysed during constant current injection at the soma that elicited a spike frequency of 20Hz with and without dendritic rectifying tonic inhibition. Membrane voltage at adjacent compartments to the soma, and all transmembrane ionic currents at the soma were recorded. Axial current flow between the soma and a dendrite (*d*) was calculated from Ohm’s law using axial resistivity (*R*_*i*_), somatic voltage (*V*_*soma*_) and dendritic voltage (*V*_*d*_). Total axial current at the soma (*I*_*Ax*_) is the sum of all axial currents:

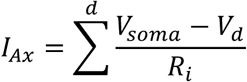

*I*_*Ax*_ was adjusted for somatic surface area and expressed as current density (**Fig. 5**). Total membrane current is defined as the sum of all ionic (*I*_*Ion*_), axial and injected current (*I*_*stim*_). When considering the current-balance equation, total membrane current is equivalent to the first derivative of the time-voltage trace adjusted by somatic membrane capacitance (*c*):

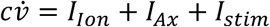

This justifies use of the derivative of the time-voltage trace to compare qualitative changes in total membrane current.

The ISI was defined as the interval between the onset of two consecutive AP’s (Section 6) during steady-state firing after a 1 second current injection. Δ total membrane current was calculated by subtracting total membrane current with and without tonic inhibition during an ISI (**Fig. 5b**). To ensure calculation of Δ total membrane current was not subject to numerical error, simulations were performed with a fixed time-step of shorter duration until differences in waveform were no longer visible by eye. Results presented use a time step of 0.001ms.

Changes in axial (Δ axial) and individual transmembrane ionic currents during an ISI were calculated using the same approach (**Fig. 5e**). To determine the contribution of axial and transmembrane ionic currents to Δ total membrane current over T_1_ and T_2_, a normalised change in membrane charge transfer was calculated. The contribution of axial charge transfer (Δ axial charge) over period T_1_ is given by:

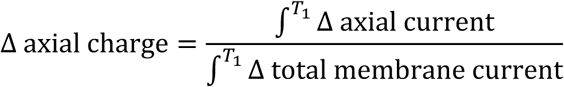

The contribution of all ionic currents over period T_1_ and T_2_ was determined in the same manner.

### 8) Simplified neuron modelling

Simplified neuron models were developed in three stages. First, channel mechanism used in detailed models were re-expressed in terms of a variable that evolves over one of three time scales: fast (*v*), slow (*w*) and ultraslow (*u*). Second, single compartment models containing these mechanisms were optimised to fit the features of a detailed bAC and cNAC model with and without tonic inhibition. Finally, optimised simple models were only analysed if the model ‘with’ tonic inhibition exhibited enhanced AP repolarisation and early recovery from AP repolarisation compared to the model ‘without’ tonic inhibition, similar to their detailed counterparts.

Channel mechanisms were simplified using a technique based on separation of timescales and equivalent voltages(60, 61). First, Na_T_ channel activation responsible for AP upstroke is considered instantaneous with respect to voltage (*v*). Using a standard Hodgkin-Huxley formulation, the Na_T_ activation variable (*m*) is therefore set to its steady-state value (*m*_*NaT,∞*_*(v)*) and Na_T_ current density (*I*_*NaT*_) given by:

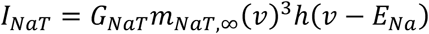

where *G*_*NaT*_ is peak Na_T_ channel conductance, *m*_*NaT,∞*_*(v)* steady-state Na_T_ conductance, *E*_*Na*_ Na reversal potential and *h*_*NaT*_ the Na_T_ channel inactivation variable.

Next, the K_P_ activation variable (*w*) was used in the bAC model to govern dynamics of other variables evolving over a slow time-scale that generate the AP downstroke, AHP and ISI: Na_T_ inactivation, Kv_3.1_ activation, Na_P_ activation and I_h_ channel activation. For the cNAC model, AP feature could only be reproduced if *w* was governed by kinetics of Kv_3.1_ activation. For the bAC model, the inverse of the K_P_ steady-state function (*w*_*∞*_*(v)*) describes an equivalent voltage (*v*_*w*_) for a given value of *w. I*_*Kp*_ and *I*_*Na*_, the latter re-expressed in terms of *v*_*w*_, is therefore given by:

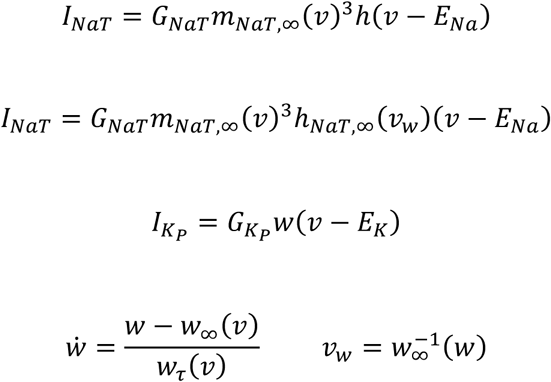

where *w*_*τ*_*(v)* is the K_P_ time constant function and *h*_*NaT,∞*_*(v*_*w*_) the Na_T_ inactivation variable steady-state function. An identical formulation was used for the simple cNAC model, with *v*_*w*_ instead given by the inverse of the Kv_3.1_ steady-state variable.

Finally, *I*_*m*_ channel activation (*u*) evolves over an ultraslow time scale and generates features of spike-frequency adaptation.

The current-balance equation for the full, simplified bAC model is given by:

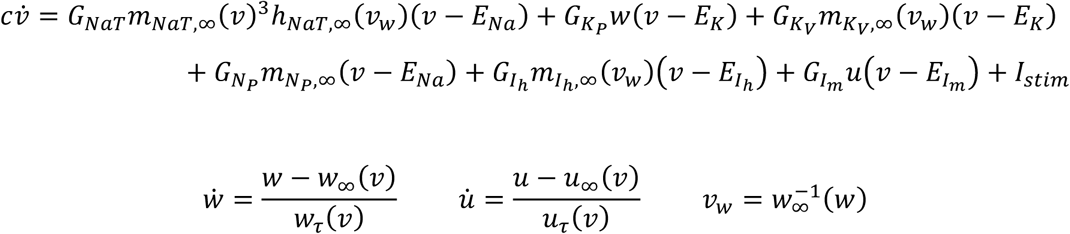

where *c* is membrane capacitance, *I*_*stim*_ input current and subscripts *N*_*P*_, *K*_*V*_ and *pas* refer to persistent Na, Kv_3.1_ and leak current respectively.

Parameters of simple models were optimised using a similar approach to Section 1. First, features used to optimise detailed models were extracted from a detailed bAC and cNAC model with and without tonic inhibition using EFEL. These features were used as means for the multi-objective function. The standard deviation for each feature was identical to those used during fitting for detailed models. For each optimised simple model, the sum of all feature errors was under 20. Ie the simplified models replicated the features of detailed models more accurately than the detailed models replicated features from *in-vitro* recordings.

Phase plane analysis of the *v-w* subsystem of simple models was performed after freezing the ultraslow variable (*u*). This approached is justified mathematically since the time constants of each variable are separated by at least an order of magnitude(61). Phase plane analysis was performed using XPPAUT(62). In **Fig. 6e** (& **Supplementary Fig. 5e**) the bottleneck is defined as a period of the trajectory that accounts for 90% of the duration of the orbit at rheobase.

### 10) Network modelling

A rate-based model developed by *Hayut et al* consisting of a population of excitatory pyramidal cells (PC) and SST interneurons was used for network simulations(36). PC and SST populations receive reciprocal synaptic connections and the PC population recurrent excitatory connections and a constant input *I*_*stim*_. Short-term synaptic plasticity uses a *Tsodyks-Markram* formulation with 3 dynamic variables: fraction of open channels (*s*), fraction of vesicles available for release (*x*) and a utilisation parameter (*u*)(63). Other synaptic parameters include conductance (*g*), initial probability of vesicle release (*U*), decay time constant of the post-synaptic current (*τ*_*s*_) and recovery time constants from facilitation and depression (*τ*_*f*_, *τ*_*r*_ respectively). Synaptic dynamics from neuronal population *j* to population *i* are as follows:

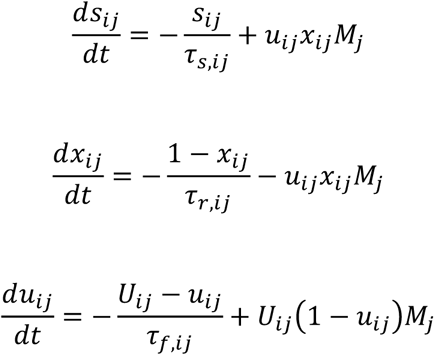

Mean firing rates (*M*) for PC and SST neuronal populations, denoted by subscripts *P* and *S*, are given by:

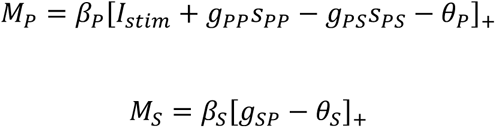

Where *θ* determines neuronal rheobase, *β* neuronal gain and []_+_ is the linear-threshold function:

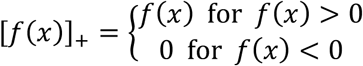

Synapses from PC to SST neurons are facilitating, and those from SST to PC neurons depressing. All parameters used in this study are identical to those in *Hayut et al* unless otherwise stated^21^.

The sensitivity analysis in **Fig. 7** was performed by simulating network activity across different *g*_*pp*_ and *I*_*stim*_ values before and after a 20% and 40% increase in *β*_*s*_. Network simulations were performed using PyDSTool(64). The network was considered to exhibit sustained gamma-frequency oscillations if oscillations persisted for over 1000 ms. Oscillations lasting between 500-1000ms were considered transient.

### 10) Statistics

All statistical analyses were performed using Python and the SciPy library. Results are presented as mean ± s.e.m. Comparison of gain values between fast spiking and non-fast spiking E-types were tested using Welch’s *t*-test, and between all Petilla E-types using one-way ANOVA with Tukey’s *post hoc* test. Gain values between fast spiking and non-fast spiking E-types from experimental recordings were tested the using the Mann-Whitney U test. We based sample sizes for our modelling results on a pilot study of one fast spiking and one non-fast spiking model. Here, a Δ gain of ∼ −10% and +10% was observed. Assuming a variance of 5% we calculated a sample of at least 8 to ensure adequate power. For experimental studies, we used a similar sample size to our models under the assumption that our models were predictive. Our sample sizes are also comparable to those reported in previous publications exploring the impact of a neuromodulator upon single neuron excitability(7, 31, 39). Differences were considered significant if \**P < 0*.*01*, \*\**P < 0*.*001*. n.s. denotes not significant.

## SUPPLEMENTARY FIGURES

**Supplementary Fig. 1.**
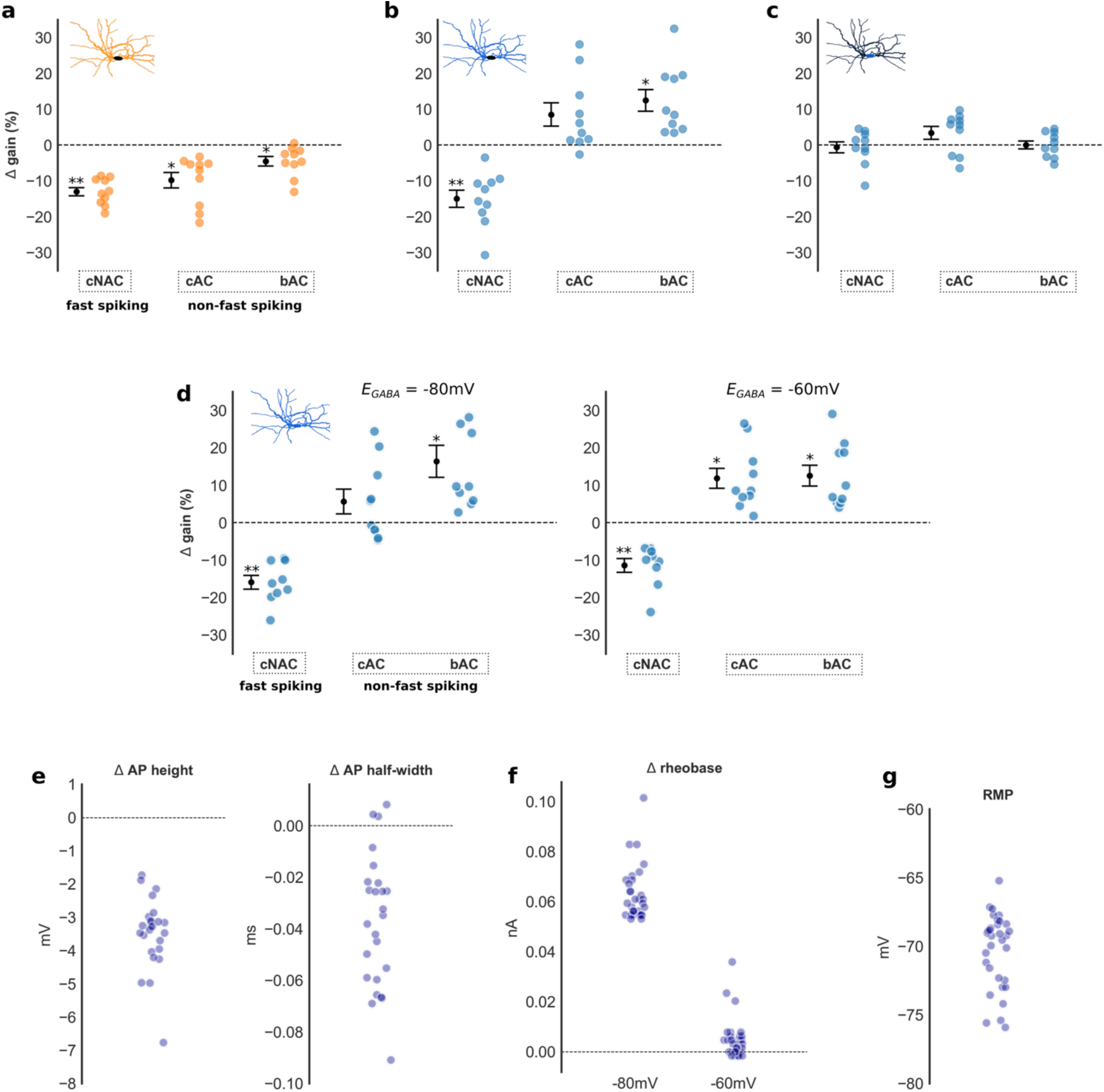
**a-c)** Δ gain in all models grouped by E-type (*E*_*GABA*_ = −80mV) for non-rectifying (**a**) and rectifying (**b**) tonic inhibition acting upon the dendritic tree, and rectifying tonic inhibition acting upon the soma (**c**). A small but significance difference in Δ gain was observed between fast spiking and non-fast spiking models with non-rectifying inhibition (−13.0% ± 1.2 vs −7.2% ± 1.4, Welch’s *t*-test, *t*(28) = −3.2, *P* < 0.01), and large difference in Δ gain with rectifying tonic inhibition (−15.0% ± 1.1 vs 10.5% ± 1.4, Welch’s *t*-test, *t*(28) = −3.4, *P* < 0.001). A significant difference in Δ gain was not observed if rectifying tonic inhibition only acted upon the soma (−0.6% ± 1.5 vs 1.7% ± 1.1, Welch’s *t*-test, *t*(28) = −1.3, n.s.). **d)** Δ gain in all models grouped by E-type for rectifying tonic inhibition acting upon all regions of the interneuron model for *E*_*GABA*_ of −80 (left) and −60mV (right). A significant difference in Δ gain was observed between fast spiking and non-fast spiking models for *E*_*GABA*_ of −80mV (−16.0% ± 1.8 vs 11.0% ± 2.9, Welch’s *t*-test, *t*(28) = −6.0, *P* < 0.001) and −60mV (−11.4% ± 1.9 vs 12.2% ± 1.9, Welch’s *t*-test, *t*(28) = −7.7, *P* < 0.001). **e)** Changes in AP features for all models in the presence of rectifying tonic inhibition. Rectifying tonic inhibition produced reductions of either AP half-width or height within every model. **f)** Changes of rheobase in all models for *E*_*GABA*_ of −80 and −60mV. **g)** Resting membrane potential (RMP) of all models. An *E*_*GABA*_ of −80 and −60mV is hyperpolarised and depolarised, respectively, relative to the RMP in every model. Results presented as mean ± s.e.m. Asterix denotes a significant Δ gain value compared to Δ gain = 0%, one-sample *t*-test, *\*P* < 0.01, *\*\*P* < 0.001.

**Supplementary Fig. 2.**
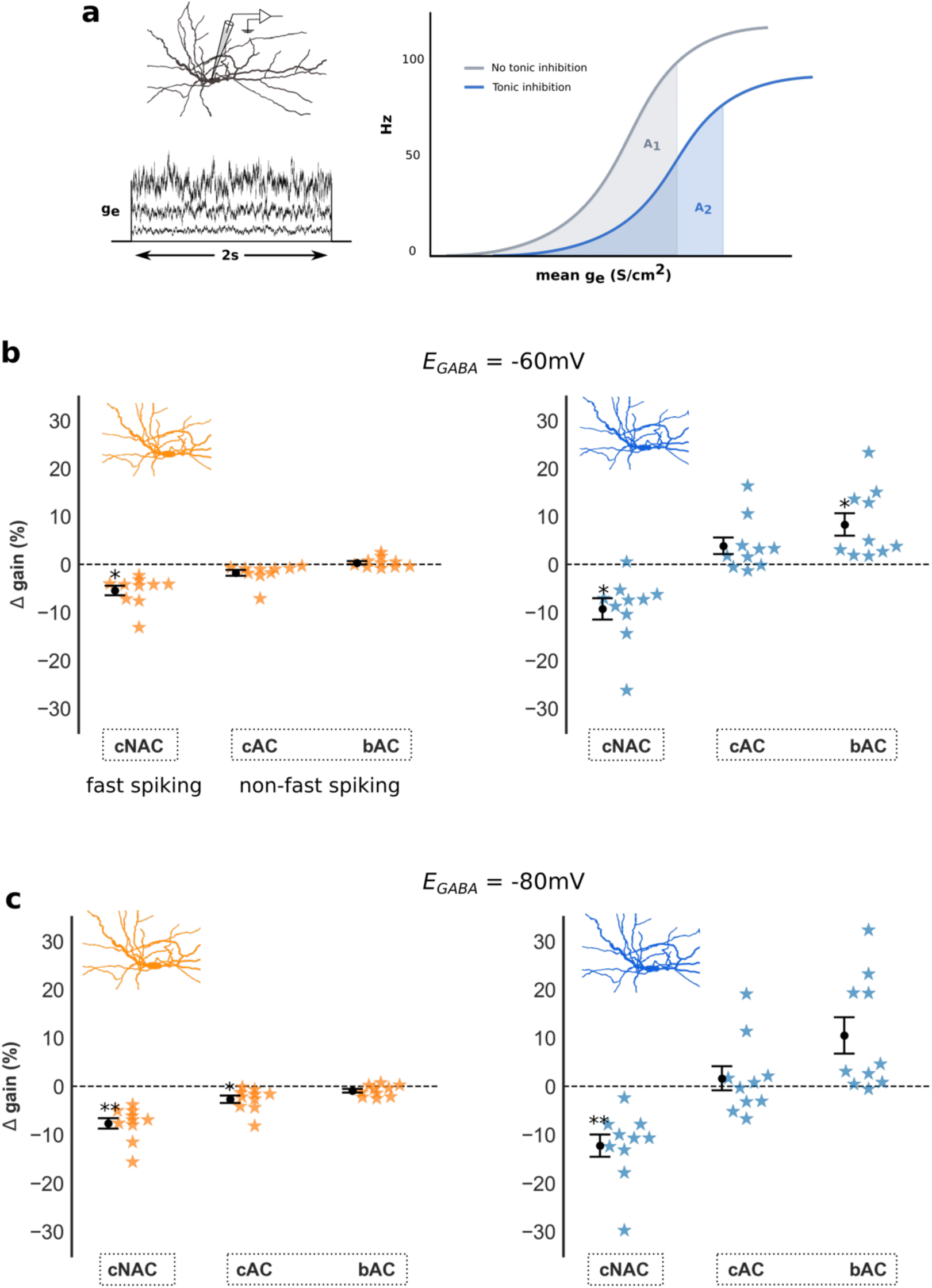
**a)** Excitatory conductance noise (*g*_*e*_) of increasing mean and variance was injected into the soma and Δ gain calculated using identical method to **Fig. 1b**. **b-c)** Δ gain in all models grouped by E-type for *E*_*GABA*_ of −60 (**b**) and −80mV (**c**) in response to noisy input with rectifying (blue) and non-rectifying (orange) tonic inhibition. Both rectifying and non-rectifying tonic inhibition induced significant differences in Δ gain between fast spiking and non-fast spiking E-types, however the magnitude of this difference was greater in the presence of rectifying tonic inhibition (−9.3% ± 2.2 vs 6.1% ± 1.5, Welch’s *t*-test, *t*(28) = −5.7, *P* < 0.001 *E*_*GABA*_ of −60mV, and −12.3% ± 2.3 vs 6.1% ± 2.4, Welch’s *t*-test, *t*(28) = −5.5, *P* < 0.001 *E*_*GABA*_ of −80mV). Asterix denotes a significant Δ gain value for each Petilla E-type compared to Δ gain = 0%, one-sample *t*-test, *\*P* < 0.01, *\*\*P* < 0.001.

**Supplementary Fig. 3.**
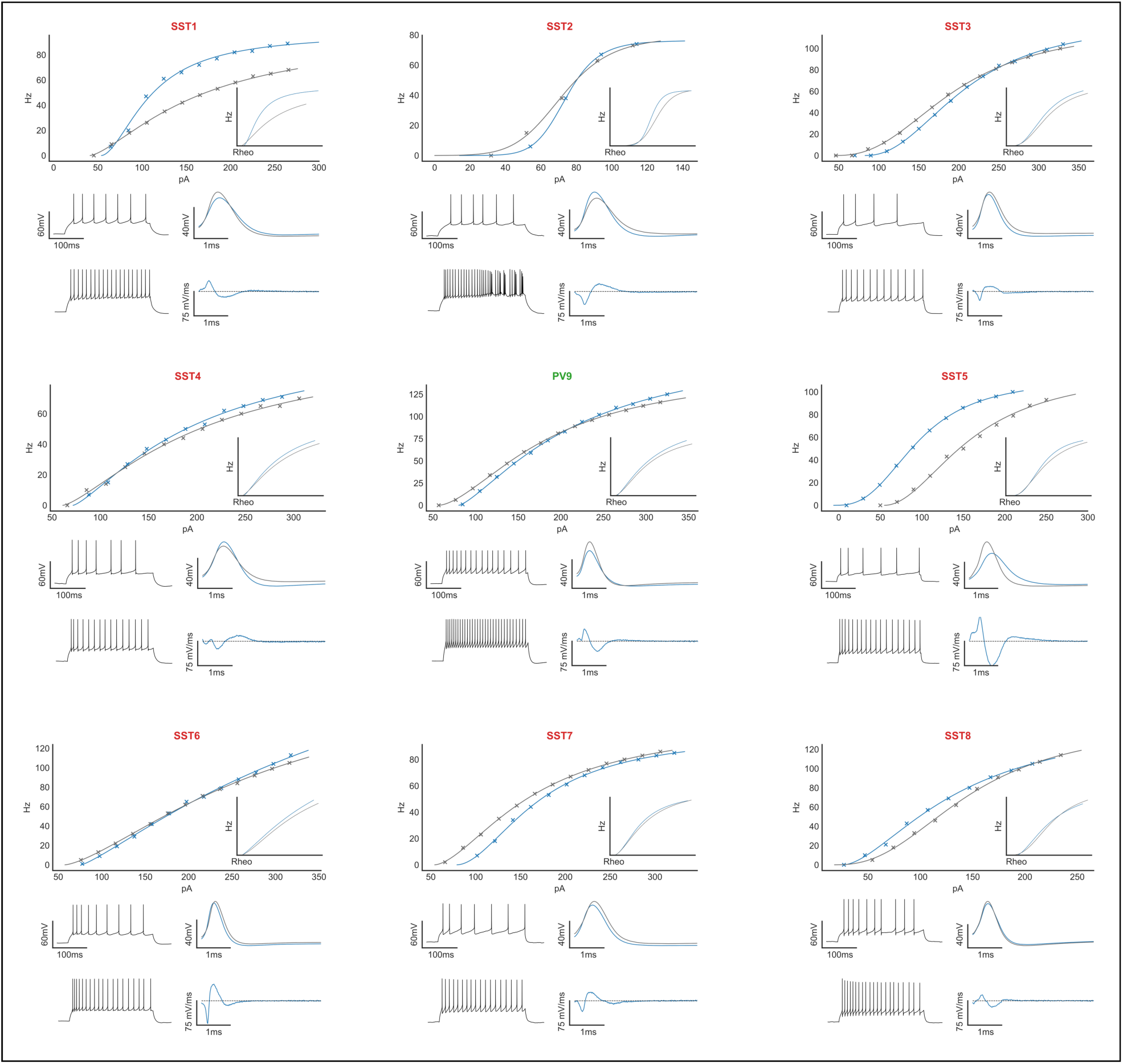
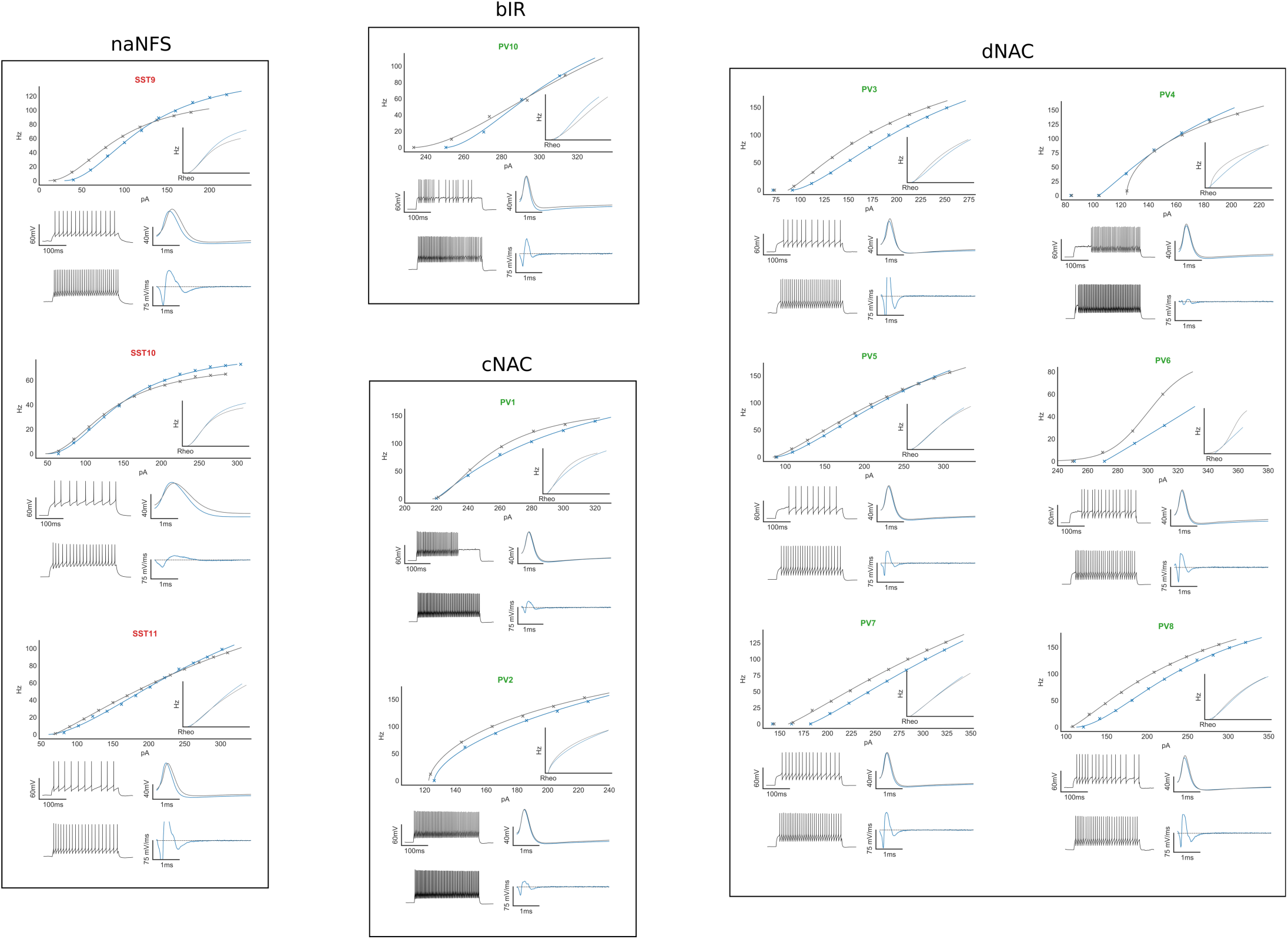
Current-frequency relationship of all recorded interneurons grouped by E-type. Inset shows current-frequency relationship adjusted for rheobase. Time-voltage traces (lower left) show electrophysiologic characteristics at rheobase (top) and 20nA above rheobase (bottom) without tonic inhibition. Lower right traces show first 4ms of time-voltage trace after AP onset during steady-state firing (top) and Δ dV/dt over the same period (bottom) providing a qualitative estimate of Δ total membrane current. Electrophysiologic features for each cell are found in **Table 1**.

**Supplementary Fig. 4.**
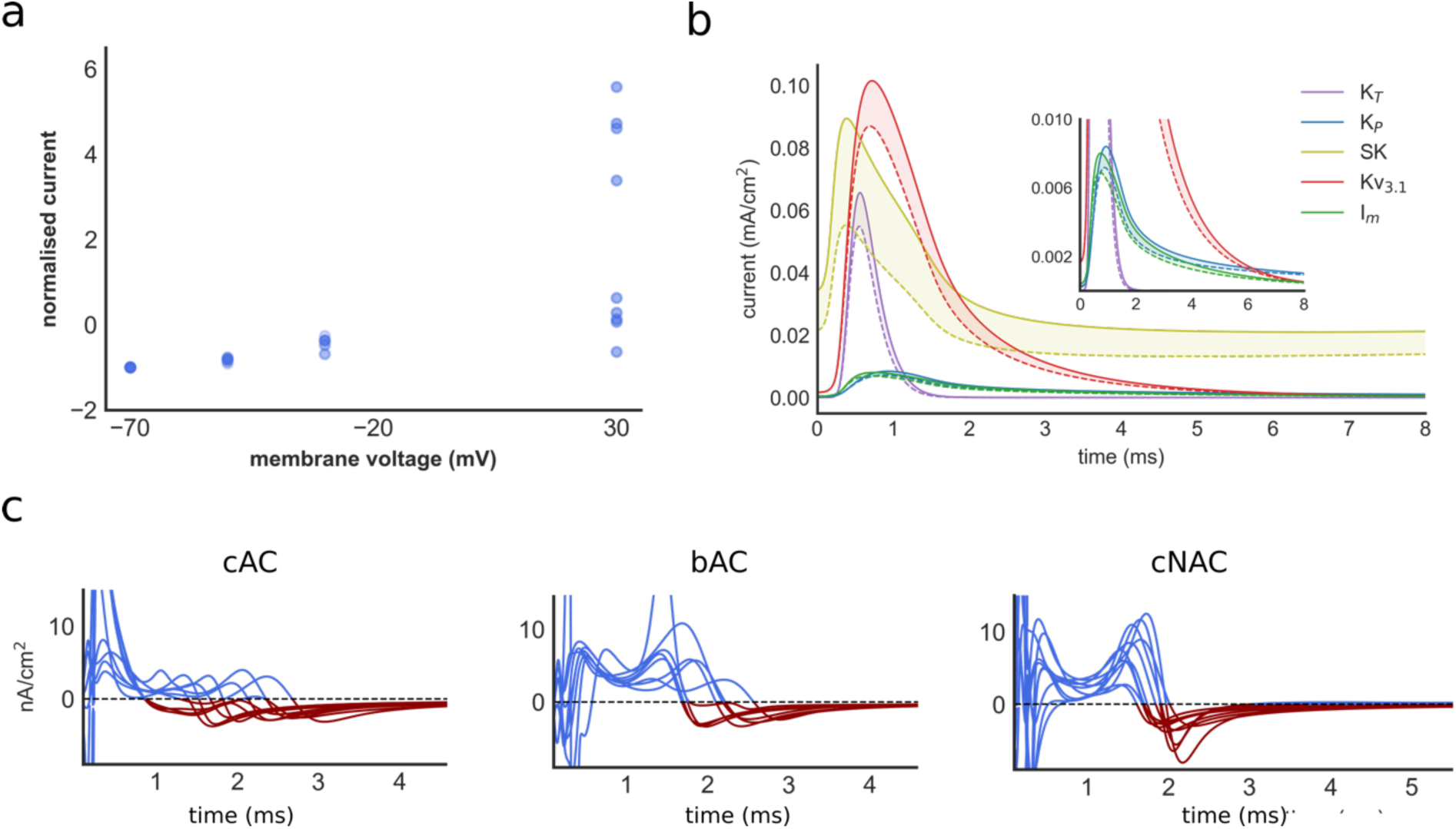
**a)** Current-voltage relationship of 9 Sst-positive interneurons. Current is normalised to holding current at −70mV. Four interneurons show marked outward current rectification. **b)** Potassium currents during an inter-spike interval in one interneuron model without (solid line) and with (dashed line) rectifying tonic inhibition. Filled region highlights the change in potassium current. The presence of tonic inhibition reduces the activation of all voltage-dependent potassium currents. **c)** Δ total membrane current, recorded at the soma in the presence of dendritic rectifying tonic inhibition, in all models grouped by E-type. Rectifying tonic inhibition enhanced AP repolarisation in all models (T_1_) and promoted earlier recovery from AP repolarisation (T_2_, depolarising change in total membrane current highlighted in red). Channel abbreviations: K_T_ = transient K, K_P_ = persistent K, SK, = Calcium-activated K

**Supplementary Fig. 5.**
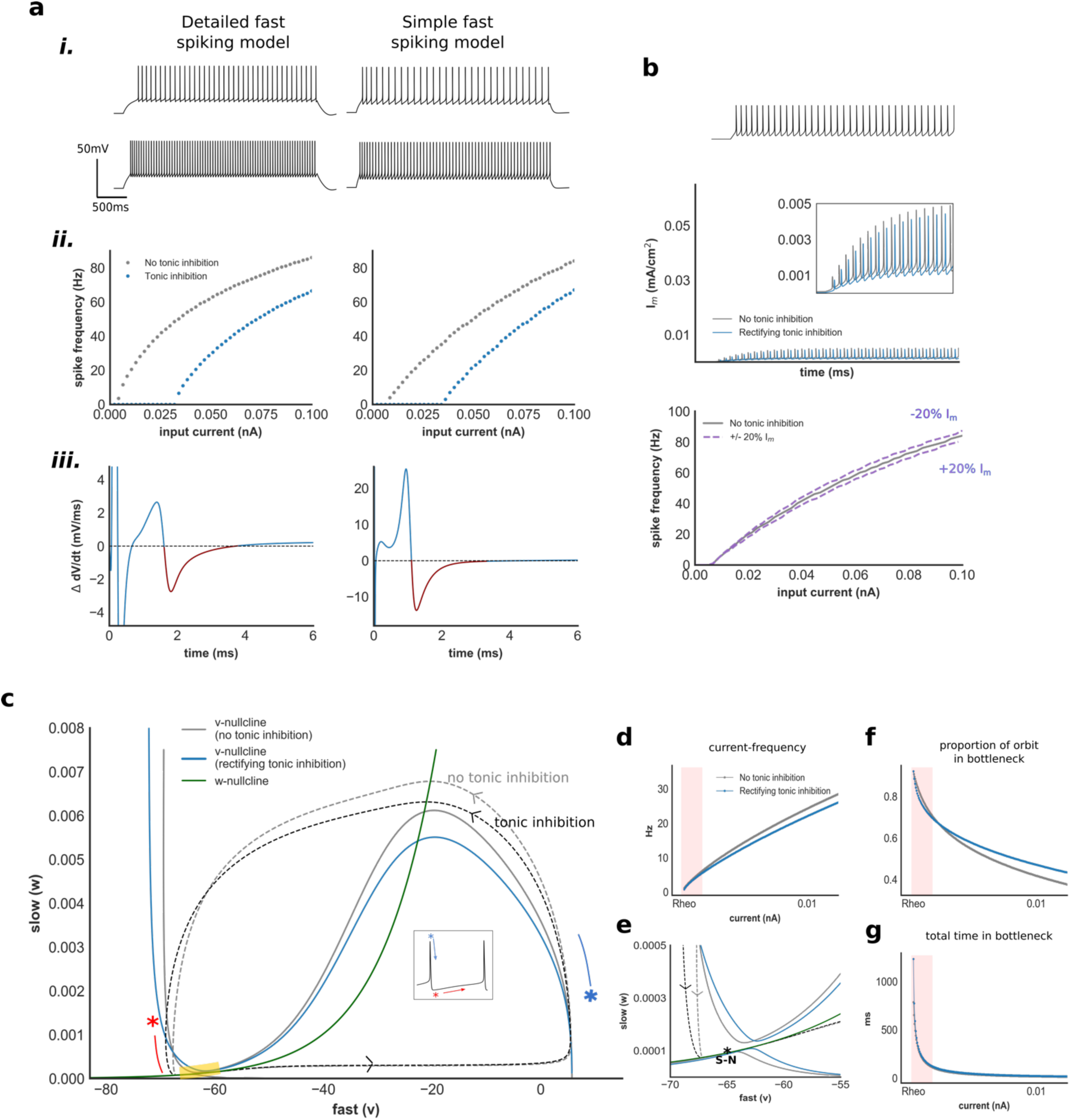
**a**) Electrophysiological features (i), current-frequency relationship (ii) and change in total membrane current (iii) within a simplified fast spiking models and its detailed counterpart. **b)** I_m_ current generated by the simple fast spiking model without (grey) and with (blue) tonic inhibition. Impact of changes in I_m_ current upon gain in the fast spiking model. **c)** Phase-plane of the *v*-*w* subsystem of the fast spiking model, and orbits during AP generation, without and with tonic inhibition. Yellow region denotes bottleneck which occupies 90% of the duration of the orbit at spike onset. **d)** Current-frequency relationship of the *v*-*w* system. **e)** Phase-plane of the *v*-*w* system at the bottleneck. The transition from rest to spiking occurs via saddle-node (S-N) bifurcation. During the AP downstroke and AHP, w deactivates rapidly compared to the fast spiking model, and the trajectory traverses the bottleneck adjacent to the *w* nullcline (c/w **Fig. 6c**). Consequently, reduced activation of *w* with tonic inhibition has minimal impact upon the trajectory through the bottleneck. **f)** Proportion of orbit spent traversing the bottleneck, and proportion of time within the bottleneck (**g**) with increasing input current. Tonic inhibition does not increase the rate of scaling (gain) through this region with increasing input current.

## REFERENCES

1. Glykys J, Mody I (2007) Activation of GABA(A) receptors: views from outside the synaptic cleft. Neuron 56(5):763–770.

2. Farrant M, Nusser Z (2005) Variations on an inhibitory theme: phasic and tonic activation of GABA(A) receptors. Nat Rev Neurosci 6(3):215–229.

3. Lee V, Maguire J (2014) The impact of tonic GABA(A) receptor-mediated inhibition on neuronal excitability varies across brain region and cell type. Front Neural Circuits 8:3.

4. Belelli D, et al. (2009) Extrasynaptic GABA(A) receptors: form, pharmacology, and function. J Neurosci 29(41):12757–12763.

5. Pavlov I, Savtchenko LP, Kullmann DM, Semyanov A, Walker MC (2009) Outwardly rectifying tonically active GABA(A) receptors in pyramidal cells modulate neuronal offset, not gain. J Neurosci 29(48):15341–15350.

6. Pavlov I, et al. (2014) Tonic GABA(A) conductance bidirectionally controls interneuron firing pattern and synchronization in the CA3 hippocampal network. Proc Natl Acad Sci U S A 111(1):504–509.

7. Song I, Savtchenko L, Semyanov A (2011) Tonic excitation or inhibition is set by GABA(A) conductance in hippocampal interneurons. Nat Commun 2:376.

8. Jin Z, et al. (2011) Insulin reduces neuronal excitability by turning on GABAAchannels that generate tonic current. PLoS One 6(1):e16188.

9. Birnir B, Everitt AB, Gage PW (1994) Characteristics of GABA(A) channels in rat dentate gyrus. J Membr Biol 142(1):93–102.

10. Semyanov A, Walker MC, Kullmann DM (2003) GABA uptake regulates cortical excitability via cell type-specific tonic inhibition. Nat Neurosci 6(5):484–490.

11. Davies PA, Hanna MC, Hales TG, Kirkness EF (1997) Insensitivity to anaesthetic agents conferred by a class of GABA(A) receptor subunit. Nature 385(6619):820–823.

12. Bianchi MT, Macdonald RL (2002) Slow phases of GABA(A) receptor desensitization: structural determinants and possible relevance for synaptic function. J Physiol 544(Pt 1):3–18.

13. Brickley SG, Mody I (2012) Extrasynaptic GABA(A) Receptors : Their Function in the CNS and Implications for Disease. Neuron 73(1):23–34.

14. Egawa K, Fukuda A (2013) Pathophysiological power of improper tonic GABA(A) conductances in mature and immature models. Front Neural Circuits 7(170). doi:10.3389/fncir.2013.00170.

15. Meltzer-Brody S, et al. (2018) Brexanolone injection in post-partum depression: two multicentre, double-blind, randomised, placebo-controlled, phase 3 trials. Lancet 392(10152):1058–1070.

16. Darmani G, et al. (2016) Effects of the Selective α5-GABA(A) Receptor Antagonist S44819 on Excitability in the Human Brain: A TMS–EMG and TMS–EEG Phase I Study. J Neurosci 36(49):12312–12320.

17. Silver RA (2010) Neuronal arithmetic. Nat Rev Neurosci 11(7):474–489.

18. Cavelier P, Hamann M, Rossi D, Mobbs P, Attwell D (2005) Tonic excitation and inhibition of neurons : ambient transmitter sources and computational consequences. Prog Biophys Mol Biol 87(1):3–16.

19. Holt GR, Koch C (1997) Shunting inhibition does not have a divisive effect on firing rates. Neural Comput 9(5):1001–1013.

20. Prescott SA, De Koninck Y (2003) Gain control of firing rate by shunting inhibition: roles of synaptic noise and dendritic saturation. Proc Natl Acad Sci U S A 100(4):2076–2081.

21. Mitchell SJ, Silver RA (2003) Shunting inhibition modulates neuronal gain during synaptic excitation. Neuron 38(3):433–445.

22. Kaila K, Price TJ, Payne JA, Puskarjov M, Voipio J (2014) Cation-chloride cotransporters in neuronal development, plasticity and disease. Nat Rev Neurosci 15(10):637–654.

23. Clemente-Perez A, et al. (2017) Distinct Thalamic Reticular Cell Types Differentially Modulate Normal and Pathological Cortical Rhythms. Cell Rep 19(10):2130–2142.

24. Hattori R, Kuchibhotla K V., Froemke RC, Komiyama T (2017) Functions and dysfunctions of neocortical inhibitory neuron subtypes. Nat Neurosci 20(9):1199–1208.

25. Kepecs A, Fishell G (2014) Interneuron cell types are fit to function. Nature 505(7483):318– 326.

26. Tremblay R, Lee S, Rudy B (2016) GABAergic Interneurons in the Neocortex: From Cellular Properties to Circuits. Neuron 91(2):260–292.

27. DeFelipe J, et al. (2013) New insights into the classification and nomenclature of cortical GABAergic interneurons. Nat Rev Neurosci 14(3):202–16.

28. Groen MR, et al. (2014) Development of dendritic tonic GABAergic inhibition regulates excitability and plasticity in CA1 pyramidal neurons. J Neurophysiol 112(2):287–299.

29. Destexhe A, Rudolph M, Fellous JM, Sejnowski TJ (2001) Fluctuating synaptic conductances recreate in vivo-like activity in neocortical neurons. Neuroscience 107(1):13–24.

30. Druckmann S, Hill S, Schürmann F, Markram H, Segev I (2013) A Hierarchical Structure of Cortical Interneuron Electrical Diversity Revealed by Automated Statistical Analysis. Cereb Cortex 23(12):2994–3006.

31. Sanchez G, Rodriguez MJ, Pomata P, Rela L, Murer MG (2011) Reduction of an Afterhyperpolarization Current Increases Excitability in Striatal Cholinergic Interneurons in Rat Parkinsonism. J Neurosci 31(17):6553–6564.

32. Hsiao CF, Trueblood PR, Levine MS, Chandler SH (1997) Multiple Effects of Serotonin on Membrane Properties of Trigeminal Motoneurons In Vitro. J Neurophysiol 77(6):2910–2924.

33. Wallén P, et al. (1989) Effects of 5-hydroxytryptamine on the afterhyperpolarization, spike frequency regulation, and oscillatory membrane properties in lamprey spinal cord neurons. J Neurophysiol 61(4):759–768.

34. Ermentrout B (1998) Linearization of F-I Curves by Adaptation. Neural Comput 10(7):1721–1729.

35. Haider B, McCormick DA (2009) Rapid Neocortical Dynamics: Cellular and Network Mechanisms. Neuron 62(2):171–189.

36. Hayut I, Fanselow EE, Connors BW, Golomb D (2011) LTS and FS inhibitory interneurons, short-term synaptic plasticity, and cortical circuit dynamics. PLoS Comput Biol 7(10):e1002248.

37. Veit J, Hakim R, Jadi MP, Sejnowski TJ, Adesnik H (2017) Cortical gamma band synchronization through somatostatin interneurons. Nat Neurosci 20(7):951–959.

38. Olah S, et al. (2009) Regulation of cortical microcircuits by unitary GABA-mediated volume transmission. Nature 461(7268):1278–1281.

39. Xiang Z, Huguenard JR, Prince DA (1998) Cholinergic switching within neocortical inhibitory networks. Science (80-) 281(5379):985–988.

40. Obermayer J, et al. (2018) Lateral inhibition by Martinotti interneurons is facilitated by cholinergic inputs in human and mouse neocortex. Nat Commun 9(1):4101.

41. Stringer C, et al. (2016) Inhibitory control of correlated intrinsic variability in cortical networks. Elife 5:e19695.

42. Wang LY, Gan L, Forsythe ID, Kaczmarek LK (1998) Contribution of the Kv3.1 potassium channel to high frequency firing in mouse auditory neurones. J Physiol 509(1):183–194.

43. Izhikevich EM (2003) Simple model of spiking neurons. IEEE Trans Neural Networks 14(6):1569–1572.

44. Izhikevich EM (2007) Dynamical Systems in Neuroscience: Geometry of Excitability and Bursting (The MIT Press).

45. Verdoorn TA, Draguhn A, Ymer S, Seeburg PH, Sakmann B (1990) Functional properties of recombinant rat GABA(A) receptors depend upon subunit composition. Neuron 4(6):919– 928.

46. Okaty BW, Miller MN, Sugino K, Hempel CM, Nelson SB (2009) Transcriptional and Electrophysiological Maturation of Neocortical Fast-Spiking GABAergic Interneurons. J Neurosci 29(21):7040–7052.

47. Ramaswamy S, et al. (2015) The neocortical microcircuit collaboration portal: a resource for rat somatosensory cortex. Front Neural Circuits 9. doi:10.3389/fncir.2015.00044.

48. Markram H, et al. (2015) Reconstruction and Simulation of Neocortical Microcircuitry. Cell 163(2):456–92.

49. Hines ML, Carnevale NT (1997) The NEURON simulation environment. Neural Comput 9(6):1179–1209.

50. Kasugai Y, et al. (2010) Quantitative localisation of synaptic and extrasynaptic GABA(A) receptor subunits on hippocampal pyramidal cells by freeze-fracture replica immunolabelling. Eur J Neurosci 32(11):1868–1888.

51. Rössert C, et al. (2016) BluePyOpt : Leveraging Open Source Software and Cloud Infrastructure to Optimise Model Parameters in Neuroscience. Front Neuroinform 10(6).

52. Druckmann S, Banitt Y, Gidon A, Sch F, Markram H (2007) A novel multiple objective optimization framework for constraining conductance-based neuron models. Front Neurosci 1(1).

53. Hay E, Hill S, Schürmann F, Markram H, Segev I (2011) Models of neocortical layer 5b pyramidal cells capturing a wide range of dendritic and perisomatic active properties. PLoS Comput Biol 7(7):e1002107.

54. Druckmann S, et al. (2011) Effective Stimuli for Constructing Reliable Neuron Models. PLOS Comput Biol 7(8):534–542.

55. Ascoli G a, et al. (2008) Petilla terminology: nomenclature of features of GABAergic interneurons of the cerebral cortex. Nat Rev Neurosci 9(7):557–568.

56. Markram H, Toledo-rodriguez M, Wang Y, Gupta A (2004) Interneurons of the neocortical inhibitory system. Nat Rev Neurosci 5(10):793–807.

57. Kvitsiani D, et al. (2013) Distinct behavioural and network correlates of two interneuron types in prefrontal cortex. Nature 498(7454):363–366.

58. Fanselow EE, Connors BW (2010) The Roles of Somatostatin-Expressing (GIN) and Fast-Spiking Inhibitory Interneurons in UP-DOWN States of Mouse Neocortex. J Neurophysiol 104(2):596–606.

59. Electrophys Feature Extraction Library Available at: https://github.com/BlueBrain/eFEL.

60. Drion G, O’Leary T, Marder E (2015) Ion channel degeneracy enables robust and tunable neuronal firing rates. Proc Natl Acad Sci 112(38):E5361–E5370.

61. Franci A, Drion G, Sepulchre R (2013) Modeling the modulation of neuronal bursting: a singularity theory approach. SIAM J Appl Dyn Syst 13(2):798–829.

62. Ermentrout B, Mahajan A (2003) Simulating, Analyzing, and Animating Dynamical Systems: A Guide to XPPAUT for Researchers and Students. Appl Mech Rev 56(4):B53.

63. Tsodyks M, Markram H (1998) Neural Networks with Dynamic Synapses. Neural Comput 835(1996):821–835.

64. Clewly R, Sherwood W, LaMar M, Guckenheimer J (2007) PyDSTool, a software environment for dynamical systems modeling. Available at: http://pydstool.sourceforge.net.

